# Live-cell imaging reveals the trade-off between target search flexibility and efficiency for Cas9 and Cas12a

**DOI:** 10.1101/2023.11.16.567366

**Authors:** Lorenzo Olivi, Cleo Bagchus, Victor Pool, Ezra Bekkering, Konstantin Speckner, Wen Wu, Koen Martens, John van der Oost, Raymond Staals, Johannes Hohlbein

## Abstract

CRISPR-Cas systems have widely been adopted as genome editing tools, with two frequently employed Cas nucleases being *Spy*Cas9 and *Lb*Cas12a. Although both nucleases use RNA guides to find and cleave target DNA sites, the two enzymes differ in terms of protospacer-adjacent motif (PAM) requirements, guide architecture and cleavage mechanism. In the last years, rational engineering led to the creation of PAM-relaxed variants *Sp*RYCas9 and imp*Lb*Cas12a to broaden the targetable DNA space. By employing their catalytically inactive variants (dCas9/dCas12a), we quantified how the protein-specific characteristics impact the target search process. To allow quantification, we fused these nucleases to the photoactivatable fluorescent protein PAmCherry2.1 and performed single-particle tracking in cells of *Escherichia coli*. From our tracking analysis, we derived kinetic parameters for each nuclease with a non-targeting RNA guide, strongly suggesting that interrogation of DNA by *Lb*dCas12a variants proceeds faster than that of *Spy*dCas9. In the presence of a targeting RNA guide, both simulations and imaging of cells confirmed that *Lb*dCas12a variants are faster and more efficient in finding a specific target site. Our work demonstrates the trade-off of relaxing PAM requirements in *Spy*dCas9 and *Lb*dCas12a using a powerful framework, which can be applied to other nucleases to quantify their DNA target search.

## 1. Introduction

Clustered regularly interspaced short palindromic repeats (CRISPR) and CRISPR-associated (Cas) proteins are bacterial adaptive immunity systems that defend the host against invading genetic elements^1–3^. After unravelling their mechanistic details, several CRISPR-Cas systems have been applied for *in vivo,* sequence-specific DNA modification in all domains of life^4,5^. Delivery of a Cas protein together with its programmable guide RNA (gRNA) allows targeting of DNA sequences (protospacer) flanked by the corresponding protospacer-adjacent motif (PAM). The broad application of these genome editing tools sparked a general interest in the biophysical characterization of the molecular mechanisms driving target search of Cas nucleases^6–12^.

The target search process of Cas nucleases proceeds in four different steps (Figure 1A). After forming a complex with a gRNA^2,13^, the Cas nuclease initiates target search with a three-dimensional diffusion in the intracellular environment (3D state). Here, the protein collides with DNA and starts moving one-dimensionally along the double helix, screening for the nuclease-specific PAM (1D state). All known DNA-targeting Cas nucleases require a specific PAM motif for effective targeting^14^. Upon recognition of a PAM, the enzyme unwinds the flanking dsDNA sequence, allowing hybridisation of the gRNA with the target strand^12^ (PAM-investigating state). Complementarity of the DNA with the gRNA, especially in the first nucleotides after the PAM (seed region), causes further DNA unwinding and complete R-loop formation. This, in turn, leads to generation of a double-stranded break by wild-type Cas nucleases^15,16^. In the case of catalytically-deficient nuclease variants (dCas), R-loop formation just leads to stable binding to the target DNA^17,18^ (target-bound state). If the DNA sequence does not match the gRNA, the nuclease is released from the DNA and resumes its three-dimensional motion.

**Figure 1.**
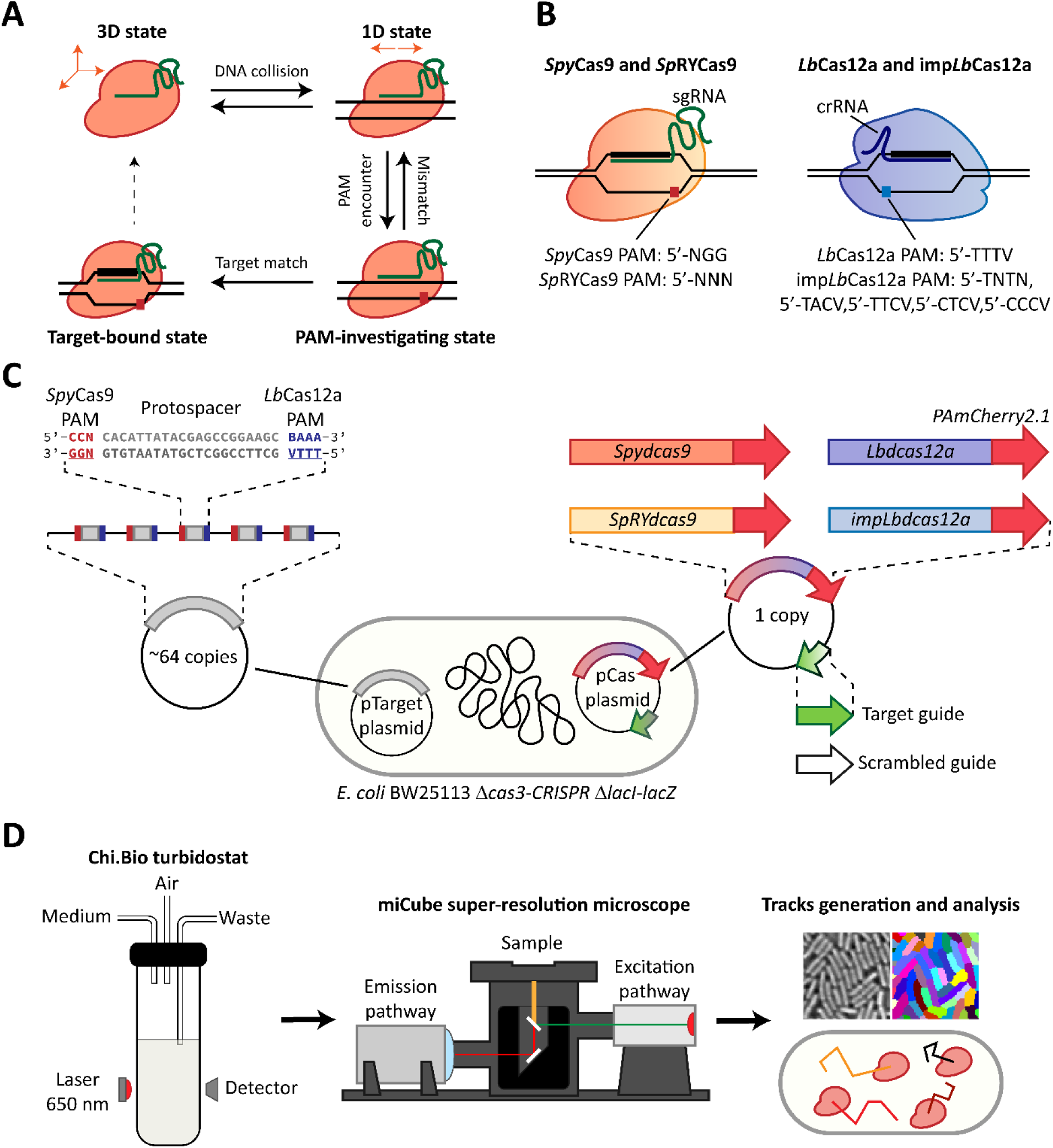
Imaging of *Spy*dCas9 and *Lb*dCas12a in live cells. **A)** The four states of Cas nucleases DNA target search, here exemplified by *Spy*dCas9. The nuclease initially diffuses three-dimensionally in the cytoplasm (state 1). Upon random collision with DNA, Cas proteins move one-dimensionally along DNA in search of a PAM (state 2). After recognising a PAM, a transient binding state is initiated, in which the nuclease attempts hybridisation of its gRNA with the DNA sequence (state 3). If the sequences do not match, the nuclease is released and reverts to state 2. If the DNA sequence and the gRNA match, a state of stable binding followed by potential DNA cleavage is achieved (state 4). **B)** Relevant characteristics of *Spy*Cas9 and *Sp*RYdCas9 (red/yellow, left), loaded with a single-guide RNA molecule (sgRNA) and recognising either a 5’-NGG-3’ or 5’-NNN-3’ PAM at the 3’ end of the protospacer, and *Lb*Cas12a and imp*Lb*dCas12a (purple/blue, right), loaded with a CRISPR RNA (crRNA) and recognising either a 5’-TTTV-3’ or a set of 5 PAMs at the 5’ end of the protospacer. **C)** Our experimental design for the sptPALM study of *Spy*dCas9, *Lb*dCas12a, *Sp*RYdCas9 and imp*Lb*dCas12a. The pCas plasmid (1 copy) harbours the Cas nuclease-PAmCherry2.1 fusion under the control of an inducible promoter and either a target or a scrambled gRNA. The pTarget plasmid (∼64 copies^37^) contains five instances of the protospacer, matching with the target guide. Each protospacer is flanked on one side by the *Spy*Cas9 PAM and on the other by the *Lb*Cas12a PAM, to allow both nucleases to recognise the same protospacer on the same pTarget plasmid. **D)** Our experimental pipeline, consisting of an initial growth phase in a turbidostat in the Chi.Bio platform, followed by sptPALM performed on the miCube platform and data analysis including cell segmentation and tracking.

In the last decade, a plethora of different CRISPR-Cas systems have been discovered and characterised. These systems are classified into 2 classes, each with 3 types and dozens of subtypes^19^. Because of their simplicity, the single-protein effector modules of type II and type V systems granted them special attention as potential genome editing tools. In particular, the type II-A *Streptococcus pyogenes* Cas9 (*Spy*Cas9) and the type V-A *Lachnospiraceae bacterium* Cas12a (*Lb*Cas12a) are widely employed nucleases for genome editing in prokaryotes and eukaryotes. Although both are single-protein effectors, the two enzymes differ in terms of their domain architecture, PAM requirements, guide architecture, as well as their DNA scanning and cleavage mechanisms(Figure 1B)^20^. *Spy*Cas9 searches for a 3 nucleotide, 5’-NGG-3’ PAM sequence via a combination of three-dimensional diffusion and unidirectional sliding, as suggested from *in vitro* experiments^11,21^. On the other hand, *Lb*Cas12a searches for a 4-nucleotide, 5’-TTTV-3’ PAM sequence using a combination of three-dimensional diffusion and hopping^12,22^. Furthermore, the *Spy*dCas9 PAM is positioned at the 3’ end of the protospacer^7^, whereas the one of *Lb*dCas12a is placed at the 5’ end^23^.

The necessity of having a PAM motif directly adjacent to the target sequence is a potential bottleneck for genome editing, as it limits the available target locations of any Cas nuclease. Thus, variants of both *Spy*Cas9 and *Lb*Cas12a have recently been engineered through rational design to considerably reduce their PAM constrains. A set of 11 amino acidic substitutions on *Spy*Cas9 led to the creation of *Sp*RYCas9, a nuclease recognising a 5’-NRN-3’ motif and minorly a 5’-NYN-3’ one, making it the first near-PAMless Cas nuclease^24^. Through a similar approach, improved *Lb*Cas12a (imp*Lb*Cas12a) was obtained by rational engineering of *Lb*Cas12a, allowing the recognition of a 5’-TNTN-3’ and four other motifs (5’-TACV-3’, 5’-TTCV-3’, 5’-CTCV-3’ and 5’-CCCV-3’)^25^.

On top of all the already collected mechanistic insights, the optimisation of genome editing efforts by CRISPR-Cas benefits from access to precise kinetic information regarding the nuclease in question. Both *Spy*Cas9^11,21,26,27^ and *Lb*Cas12a^12,22,28–30^ have been characterised at the single-molecule level *in vitro*. However, cellular environments such as the bacterial cytoplasm are much more complex and mostly occupied by DNA^31^. This cellular DNA harbours an abundance of PAM sequences, yet it usually only contains one or few targets. Thus, the flexibility of Cas nucleases, derived from the need to investigate every PAM, largely complicates target search. Initial attempts of characterising *Spy*Cas9 dynamics in absence of a protospacer reported a DNA-interaction time upper limit of 750 ms in human cell lines^8^ and of 30 ms in *Escherichia coli*^32^. Follow up single-particle tracking photo-activated localisation microscopy (sptPALM) in the bacterium *Lactococcus lactis* showed an interaction time of *Spy*dCas9 of 17 ± 4 ms with non-matching sequences and of ∼96 s when a correct protospacer was found^9^. Furthermore, a type I Cascade system was also characterised in the native host *E. coli* by sptPALM, with similar non-target interaction times of tens of milliseconds^10^. Target search dynamics of *Lb*Cas12a, on the other hand, have not yet been reported in live cells. Moreover, it is currently unknown, how different nucleases compare when searching for the same DNA target sequence and how different PAM requirements affect the process.

Here, we performed sptPALM to study the target search dynamics of the inactivated, endonuclease-deficient variants of *Spy*Cas9 (*Spy*dCas9) and *Lb*Cas12a (*Lb*dCas12a) in the model bacterium *E. coli* by fusing them to a mutant of the photo-activatable fluorophore mCherry2 (PAmCherry2.1)^33^. To further investigate the dependency of such dynamics on the PAM requirements, we added variants of the two nucleases with relaxed PAM requirements, *Sp*RYdCas9 and imp*Lb*dCas12a. We designed our system to allow all four nucleases to recognise the same protospacer (Figure 1C). The different nucleases were expressed from a single-copy number plasmid (pCas) under the control of an inducible promoter, while five instances of the protospacers were harboured on a second, high-copy number plasmid (pTarget). We obtained cultures of cells in steady-state exponential phase and then subjected them to sptPALM on the miCube open microscopy framework^9^. Due to the relatively low average track lengths of PAmCherry2.1 (∼3 steps), we analysed the resulting tracks using Monte Carlo-based diffusion distribution analysis (MC-DDA) software to extract kinetic rates^9,34^. To this end, we expanded MC-DDA to adapt to our newly proposed four-state model of Cas nuclease action. In this way, we estimated kinetic parameters governing the target search process of *Spy*dCas9, *Lb*dCas12a and their PAM-relaxed variants and quantified the duration of their non-target DNA interactions. Using the obtained kinetic parameters, we simulated the target search process of the four nucleases to establish how fast they can identify a single target present in the genome of *E. coli*. Providing targeting guides to the different nucleases then allowed an estimation of the fraction of them that were bound to their respective targets. Our work identifies potential limiting steps in PAM recognition for both *Spy*Cas9 and *Lb*Cas12 nucleases and reveals the trade-off of relaxing their PAM requirements during target search in a complex, live-cell environment. Moreover, it provides a powerful framework that might be applied to other nucleases to quantify their DNA target search dynamics.

## 2. Materials and methods

### Bacterial strains, media, and buffer preparation

All relevant bacterial strains and relative genotypes used in this study are described in Table S1. *E. coli* DH5α (NEB) was used for general cloning purposes. *E. coli* BW25113 Δ*cas3*-CRISPR1, Δ*lacIZ*^35^, henceforth referred to as *E. coli* PAM-SCANR, was used as final strain to investigate target search of Cas nucleases. All bacterial strains were stored at -80 °C.

All *E. coli* strains were routinely grown at 37 °C and 150 rpm in LB liquid medium (10 g/L tryptone, 10 g/L NaCl, 5 g/L yeast extract) for transformation purposes. M9CG/Liquid medium (5 g/L KH_2_PO_4_, 0.5 g/L NaCl, 6.78 g/L Na_2_HPO_4_, 1 g/L NH_4_Cl, 4.98 mg/L FeCl_3_, 0.84 mg/L ZnCl_2_, 0.13 mg/L CuCl_2_ · 2H_2_O, 0.1 mg/L CoCl_2_ · 6H_2_O, 0.1 mg/L H_3_BO_3_, 0.016 mg/L MnCl_2_ · 4H_2_O, 500 mg/L MgSO_4_ · 7H_2_O, 11 mg/L CaCl_2_, 3% V/V glycerol, 2 g/L casamino acids) was used during single-particle tracking experiments. LB medium was eventually supplemented with agar (1.5% w/V) to obtain solid media and antibiotics for plasmid propagation (17 mg/L chloramphenicol, 50 mg/L kanamycin). M9CG medium was eventually supplemented with proper antibiotics for plasmid propagation (17 mg/L chloramphenicol, 50 mg/L kanamycin), with anhydrotetracycline (aTc) to induce expression of Cas nucleases and with agarose (1.5% w/V final concentration) to immobilise cells for single-particle tracking experiments.

Phosphate-buffered saline solution (PBS) (NaCl 8 g/L, KCl 0.2 g/L, Na_2_HPO_4_ 1.42 g/L, KH_2_PO_4_ 0.24 g/L) was used to wash cells during sample preparation for single-particle tracking experiments.

### DNA manipulation and transformation

For cloning purposes, DNA fragments were amplified through PCR using Q5® High-Fidelity DNA Polymerase (NEB) following manufacturer instructions. Specific oligonucleotides and DNA fragments were designed and synthesised (IDT). Amplicons were isolated and purified using Zymoclean Gel DNA Recovery Kits (Zymo Research). Assembly was performed using NEBuilder® HiFi DNA Assembly Master Mix (NEB). Plasmid DNA was isolated and purified using GeneJET Plasmid Miniprep kit (Thermo Fisher Scientific). All plasmids were confirmed by sequencing through Oxford Nanopore (Plasmidsaurus). Confirmed plasmids were stored at -20 °C until use.

All *E. coli* cells were made chemically competent in house^36^. Chemically competent cells were transformed with plasmids through heat shock.

### Design and construction of the pCas-PAmCherry2.1 plasmid series

The pCas-PAmCherry2.1 plasmid series was based off the pBeloBAC11 plasmid^37^ and contained a codon-optimised sequence of either *Spy*dCas9 or *Lb*dCas12a. Each nuclease was fused through a flexible linker (GSGSS) to a mutant of PAmCherry2 containing an aminoacidic substitution that replaced a methionine with a leucine in position 10. This mutation was introduced to abolish a reported internal translation initiation site^38^ and generated PAmCherry2.1. The mutation was introduced by site-directed mutagenesis with primers BG27593 and BG27594. In addition, each plasmid contained either a targeting or a scrambled guide RNA for the corresponding deactivated nuclease. The deactivated nuclease gene was placed under the control of the tetracycline-inducible promoter PL-TetO1 and directly downstream of a bicistronic design^39^. The guide RNA was placed under the control of the constitutive promoter pJ23119. All plasmids used in this study are described in Table S2. The sequence of all oligonucleotides and DNA fragments used to introduce mentioned mutations is available in Table S3.

### Design and construction of pCas-PAmCherry2.1 plasmids carrying PAM-relaxed nucleases

The pCas-PAmCherry2.1 plasmids previously constructed were modified by introducing appropriate mutations to the deactivated nucleases to obtain their PAM-relaxed variants. For *Sp*RYdCas9^24^, a DNA fragment containing mutations to obtain the amino acidic substitutions L1111R, D1135L, S1136W, G1218K, E1219Q, N1317R, A1322R, R1333P, R1335Q and T1337R was synthesised (BG30037). Primers BG30042 and BG30043 were used to amplify the remainder of the *Spy*dCas9 sequence and introduce a mutation causing the A61R amino acidic substitution. The pBeloBAC11 backbone, the PL-TetO1 promoter, the bicistronic design, the PAmCherry2.1 sequence and either the targeting or scrambled guide were amplified from either p*Spy*dCas9-PAmCherry2.1_Target or p*Spy*dCas9-PAmCherry2.1_Scrambled with primers pairs BG30039 and BG30041 and BG30038 and BG30040. Fragments were then assembled to yield plasmids p*Sp*RYdCas9-PAmCherry2.1_Target and p*Sp*RYdCas9-PAmCherry2.1_Scrambled.

For imp*Lb*dCas12a^25^ a similar approach was followed, with mutations encoding substitutions G532R, K538V, Y542R and K595R contained in a synthetic DNA fragment (BG29090) and substitution D156R introduced through primers BG29088 and BG29089. The remainder of p*Lb*dCas12a-PAmCherry2.1_Target or p*Lb*dCas12a-PAmCherry2.1_Scrambled was amplified with primer pairs BG29091 and BG30039 and BG30040 and BG29087. Fragments were assembled to yield plasmids pimp*Lb*dCas12a-PAmCherry2.1_Target and pimp*Lb*dCas12a-PAmCherry2.1_Scrambled. All plasmids used in this study are described in Table S2. The sequences of all oligonucleotides and DNA fragments used to introduce mentioned mutations are available in Table S3.

### Design and construction of the pTarget plasmid

The pTarget plasmid was based on a member of a previously reported pSC101 library^40^. The member of the pSC101 library used contained a K102E Rep101 mutation (pSC101-K102E), leading to an average copy number of 64^40^. The pTarget plasmid harboured a locus containing 5 instances of a protospacer flanked by PAM motifs for both *Spy*dCas9 (5’-NGG) and *Lb*dCas12a (5’-TTTV), interspaced by 30 bp of random sequence. To obtain this locus, five sets of complementary oligonucleotides were designed, each containing the protospacer module at its centre, flanked at each side by 11 bp of random DNA and a unique 4 bp motif to use as overhang. Each set of oligonucleotides was mixed (200 pmol of each), the temperature was increased to 95 °C and then slowly lowered to 12 °C at 0.05 °C/sec to allow annealing. BbsI sites were introduced in pSC101-K102E via PCR and the resulting fragment was mixed with the 5 dsDNA fragments containing protospacers. The mixture was incubated with BbsI and T4 ligase at 37 °C for 2 h to yield the pTarget plasmid. All plasmids used in this study are described in Table S2.

### Turbidostat cultivation and growth rate assessment

The *E. coli* strains to image were grown overnight in LB medium supplemented with kanamycin and chloramphenicol at 37 °C, 180 rpm to an OD_600nm_ of ∼4. The densely-grown culture was then used to start a turbidostat cultivation using the Chi.Bio platform^41^. Experiments were started according to manufacturer instructions for fluidic lines, electrical connections and user interface. Chambers were filled with 20 mL of M9CG supplemented with kanamycin and chloramphenicol into which the overnight culture was inoculated to an initial OD_600nm_ of 0.05. Cultivation temperature was set to 37 °C and default settings were employed for stirring and frequency of optical density measurements. A 650 nm laser was used to monitor the optical density of the culture instead of visible light to prevent accidental photoactivation of PAmCherry2.1 due to the large emission spectrum of the LED mounted on the Chi.Bio platform. The target optical density was set to 0.9 AU (corresponding to an OD_600nm_ of ∼0.4) and the media reservoir was filled with M9CG supplemented with kanamycin, chloramphenicol and aTc. To induce expression of *Spy*dCas9 and *Lb*dCas12a, we used 0.25 ng/mL of aTc and 0.15 ng/mL of aTc, respectively. The same concentration of aTc for induction used for the wild type *Spy*dCas9 and *Lb*dCas12a was also used for their PAM-relaxed variants. Cells were grown at the target optical density for at least 10 generations (∼ 10 hours) before being harvested for sptPALM.

Growth rate was estimated from the volume of medium refreshed at the steady-state. The total spent volume was divided by the vessel volume (20 mL) to estimate the number of times the population doubled and the time spent at the steady state was used to obtain the growth rate (min^-1^) and doubling times (min).

### Sample preparation and single-particle tracking experiments

To perform sptPALM of Cas nucleases in live *E. coli,* cells were collected from turbidostat in a 50 mL Falcon tube and washed three times in PBS. Supernatant was then removed, the pellet resuspended in 100 μL of PBS and 1-2 μL of the suspension was placed on M9CG agarose pads between two heat-treated glass coverslips (#1.5H, 170 mm thickness). Glasses had been previously heated at 500 °C for 30 min in a muffle furnace to remove organic impurities.

All sptPALM experiments were performed at room temperature using the miCube open microscopy framework^9^. Briefly, the microscope mounted an Omicron laser engine, a Nikon TIRF objective (100x, oil immersion, 1.49 NA, HP, SR) and an Andor Zyla 4.2 PLUS camera running at a 10 Hz for brightfield imaging acquisition and 100 Hz for sptPALM. For each imaging experiment, 300 frames were acquired at 100 ms intervals with brightfield illumination using a commercial LED light. For sptPALM, individual videos of 30000 frames were acquired at 10 ms intervals with stroboscopic illumination. Multiple videos were collected for each field of view, until exhaustion of fluorophores. A 561 nm laser with ∼0.12 W/cm^2^ power output was used for HiLo-to-TIRF illumination with 4 ms stroboscopic illumination in the middle of a 10 ms frame. 405 nm laser light was provided with stroboscopic illumination to activate PAmCherry2.1. The 405 nm laser was initially provided with low-power (μW/cm^2^ range) and with a 0.5 ms stroboscopic illumination at the beginning of a 10 ms frame^42^. Both the power and the stroboscopic illumination were progressively increased throughout the imaging until exhaustion of fluorophores. Raw data was acquired using the open source Micro-Manager software^43^. During acquisition, a 2x2 pixel binning was used, yielding an effective pixel size of 119x119 nm. The field of view was restricted to regions of interest of 256x256 pixels or smaller. The first 500 frames of each sptPALM video were discarded, to prevent attempted localisation of overlapping fluorophores and pre-bleach fluorescent contaminations in and around the cells.

### Localisation of fluorophores and cell segmentations

Fluorophores localisation and cell segmentation were performed as previously described^9^. Briefly, watershed-based segmentation^44^ (http://imagej.net/Interactive_Watershed) was performed to obtain pixel-accurate cell outlines from the brightfield images. Single-particle localisation was performed via the ImageJ^45^/Fiji^46^ plugin ThunderSTORM^47^, with added maximum likelihood estimation-based single-molecule localisation algorithms. First, a 50-frame temporal median filter (https://github.com/marcelocordeiro/medianfilter-imagej) was applied to the sptPALM movies to correct background intensity^48^. Image filtering was performed through a difference-of-Gaussians filter (Sigma1 = 2 px, Sigma2 = 8 px). The approximate localisation of molecules was determined via a local maximum with peak intensity threshold of *std*(𝑊ave. 𝐹1) 1.2 and 8-neighbourhood connectivity. Sub-pixel localisation was performed through Gaussian-based maximum likelihood estimation, with a fit radius of 4 pixels (Sigma = 1.5 px). A custom-written, MATLAB-based pipeline was used to process and analyse the imaging data (https://github.com/HohlbeinLab/2021_sptPALM_Matlab, currently restricted access).

Different output files from ThunderSTORM were combined when multiple videos had to be recorded for the same field of view. Localisations were then assigned a cell ID if they fell inside a cell and were discarded if not. Single, valid localisations were linked into tracks according to spatial and temporal distances. The tracking procedure was performed as previously reported^9^ and yielded a number of tracks present in each cell and an overall apparent diffusion coefficient distribution. For each track of up to 9 localisations, the apparent diffusion coefficient *D*\* was obtained by calculating the mean square displacement between the first *n* steps and taking the average of that, where *n* is the number of localisations minus one. The diffusion coefficients of all tracks were then collected into 85 logarithmic-divided bins from *D*\* = 0.04 μm^2^/s to *D*\* = 10 μm^2^/s. To investigate the effect on copy number on target search in presence of a protospacer, the data originating from each strain was divided into smaller sets according to copy number ranges.

### Monte-Carlo diffusion distribution analysis of Cas nucleases with a scrambled guide

To interpret the distribution of apparent diffusion coefficients obtained from Cas nucleases provided with a scrambled guide, a set number of each nuclease was simulated moving in a three linear state model (50000 for the fit, 250000 for visualising the fit) in a Monte-Carlo diffusion distribution analysis (MC-DDA). The proteins were then fitted with a general Levenberg-Marquardt procedure in MATLAB and the error was determined via a general bootstrapping approach^9^. In our analysis pipeline, each protein is assigned an initial 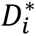 for each of its states 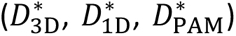 (Table S1, Supplementary Material) and an initial rate *k_i_* governing the transition between each state (*k*_3D→1D_, *k*_1D→3D_, *k*_1D→PAM_, *k*_PAM→1D_). The proteins are randomly placed inside a cell, simulated as a cylinder (length 2 μm and radius 0.5 μm) with two hemispheres at its poles (radius 0.5 μm). Each protein is randomly put in one of the three different states, based on the respective kinetic rates (see Supplementary Material). From there, the proteins are given a time before they are changed to a different state, defined as state-change time *t*_Change_ and calculated as 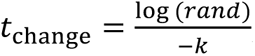, where *rand* is an evenly distributed random number and *k* is the kinetic rate governing the transition between the current and the next state. The movement of each protein is simulated with over-sampling with regards to the frame-time (*t*_step_ = 0.1 ms). In their 3D state, each nuclease moves for a distance (*s*) equal to a randomly sampled normal distribution, centred around 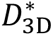 and with 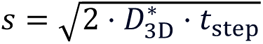 for every dimension. At every step, the *t*_change_ is subtracted with the *t*_step_. If the value becomes ≤ 0, the protein switches state and new diffusion coefficient and t_change_ are assigned. Every 10 ms after an initial equilibrium time of 200 ms, the current location of the proteins is convoluted with a random localisation error, taken from a randomly sampled normal distribution with a localisation uncertainty of σ = 0.035 μm. The simulated proteins have different numbers of localisations, leading to simulated tracks ranging from 1 to 8 steps. The amount of track of each length follows an exponential decay with a mean track length of 3 steps.

### Estimation of target finding times

As a first comparison between the several Cas nucleases, we estimated the average time needed for either 1 or 10 nucleases to find a single target in the genome of *E. coli* harbouring our plasmids. To this end, we adapted our previously developed Monte-Carlo simulation software^9^ to the four state linear model presented here. Values such as the genome size, GC content, average genome copy number (estimated at 2.3 in *E. coli* from an average doubling time of 40 min^49^) and pTarget size and copy numbers were kept constant in all cases. For each Cas nuclease to simulate, the specific kinetic rates estimated from the scrambled guide fitting were provided, together with the specific PAM sequence, the number of units per cell and the available time to find a target. We also provided a value for the probability of dissociation (*P*_dis_) while diffusing in one dimension in search of a PAM, dependent on the one-dimensional sliding length (Supplementary Information). The specific *P*_dis_ used (*P*_dis_ = 0.144) was estimated given the average 3 bp sliding length of *Spy*dCas9^11^ and was kept constant for all the nucleases. Each unit was simulated one by one. The nuclease was initialised in its 3D state, to simulate target search from *de novo* expression. From there, the proteins were assigned a *t*_Change_ value and moved between different states as mentioned before (Monte-Carlo diffusion distribution analysis of Cas nucleases with a scrambled guide). While in the 1D state, the nuclease was given a chance to encounter a PAM followed by the correct protospacer, defined as 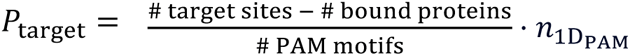. Here, 𝑛_1DPAM_ is the average number of PAM motifs encountered when sliding on a random stretch of DNA. An extensive explanation of how this value and the *P*_dis_ are estimated is available as Supplementary Information (Variables used in the estimation of target search time of Cas nucleases), together with a list of all the values used for any relevant variable (Table S2). When a nuclease encountered a PAM followed by the correct protospacer, it was assigned a *k*_Targetà3D_ rate of 0 and it was considered as irreversibly bound. A total of 2,000 cells were simulated, each containing the specified number of units. The whole simulation was then repeated 10 times and the final output was the average percentage of cells in which all the target sites were found and bound by the nuclease, together with the relative standard deviation.

The time needed for each nuclease to find its target with 50% probability (*t*_50%_) at different GC content was obtained simulating one unit of each nuclease in the same described above. Kinetic rates for each nuclease were left unchanged, together with genome size and copy number, pTarget size and copy number, *P*_dis_. The GC content was varied from 30% to 70% in steps of 10%.

### Monte-Carlo simulation of Cas nucleases with a targeting guide

To obtain target-bound fractions of Cas nucleases equipped with a targeting guide, the imaged cells were divided into smaller groups according to the number of tracks they contained. A set number of each nuclease was then simulated (250,000 for the fit, 500,000 for visualising the fit) in two different species. The first species was moving linearly in a 3 linear state model, using diffusion coefficients 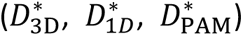 and kinetic rates (*k*_3D→1D_, *k*_1D→3D_, *k*_1D→PAM_, *k*_PAM→1D_) identified from fitting the scrambled guide distributions. The second species was simulated as static and moving with a 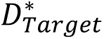 of 0.21 ± 0.05 μm^2^/s, representing the target-bound fraction. The simulation parameters were identical as the ones used for interpreting the scrambled guide diffusion coefficient distributions. The relative ratio of the two species was altered until fitting the experimental histograms for every group of cells and was judged by visual inspection. The target-bound fraction was then obtained as the number of particles in the static state over the total number of particles simulated.

## 3. Results

### Extraction of kinetic rates for *Spy*Cas9 and *Lb*Cas12a variants

To examine target search dynamics of *Spy*dCas9, *Lb*dCas12a or their PAM-relaxed variants (*Sp*RYdCas9 and imp*Lb*dCas12a), we expressed the different nuclease and tracked them in live *E. coli* cells. The nucleases were fused at their C-terminal end with the photoactivatable fluorescent protein PAmCherry2.1. This mutant variant of PamCherry2, harbouring a M10L substitution, abolishes a recently identified internal translation initiation site^38^. The same plasmid harboured either a targeting or a scrambled (non-targeting) gRNA, with or without a fully matching protospacer in the pTarget plasmid, respectively. Due to previous reports of guide-free DNA damage^50,51^, we decided that testing the behaviour of the Cas nucleases without a gRNA would constitute an additional condition outside of the scope of this study. Cells always harboured our pTarget plasmid independently from the expressed gRNA, to normalise plasmid burden throughout the tested conditions. Next, we refined our experimental pipeline. Normally, sptPALM experiments in which expression of the protein of interest is controlled by an inducible promoter requires long incubation times. This incubation interval provides sufficient time to the cells to express enough proteins and for the chromophore of the fused fluorescent proteins to mature, hence allowing enough molecules to be imaged. This potentially leads to samples with cells in stationary phase, where the nucleoid structure is more compact^52^ and core cellular processes such as replication and division are almost halted^53^. In our pipeline (Figure 1D), we strived to image live cells as close as possible to their full complexity, while avoiding set ups that would considerably increase the experimental complexity (e.g., microfluidic cultivation chambers). Therefore, *E. coli* cells harbouring our plasmids of interest were cultured in constant density settings (i.e., turbidostat) using the Chi.Bio platform^41^.

To assess the behaviour of the four tested nucleases, we created diffusional histograms belonging to cells containing the pTarget plasmid and one of the different pCas plasmids, either with a scrambled or a targeting guide RNA. Due to the temporal averaging of the diffusion coefficient in tracks that are rapidly transitioning, we fitted the distributions to obtain kinetic rates. We initially considered to treat each of the Cas states (3D, 1D, PAM-interacting, target-bound, Figure 1A) as a separate population^54,55^ and fit their diffusion coefficient distribution through an analytical expression^34,56^. However, this methods implies that all the different states are at an equilibrium and the transition between them can only be assessed thanks to rare, long-lived tracks (> 250 ms)^54^. Therefore, to grasp the full dynamics of the Cas nucleases, we decided to simulate over 50,000 particles moving in a four linear states model (Figure 1A) and used MC-DDA to fit our distributions. Each nuclease was assigned a different apparent diffusion coefficient 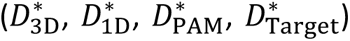, and we used a single value for localisation uncertainty (σ = 0.035 μm, or 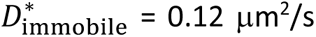). All nucleases start in a three-dimensional diffusing state (3D state), in which diffusion is solely governed by Brownian motion. Because of this, we assigned a 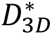 value of 2 μm^2^/s for *Spy*dCas9 and one of 2.2 μm^2^/s for *Lb*dCas12a. These values were based on the proteins hydrodynamic radius, taking into account the high viscosity in the cytoplasm^57^. After randomly encountering DNA, the nucleases transition from the 3D state to one-dimensional diffusion (1D state), where the Cas protein moves along the DNA in search of a PAM motif. *Spy*Cas9 and *Lb*Cas12a differ considerably in their mode of 1D motion. For *Spy*dCas9, this was reported to be a sliding process along the DNA^11,21^, while *Lb*dCas12a searches for its PAM through intermittent contact^12,28^ (hopping mechanism). In both cases, when moving unidirectionally on the DNA, the nucleases are apparently immobile, as they cover distances considerably lower than our localisation uncertainty (see Supplementary Material). Once a nuclease encounters a PAM, it resides on the DNA for the time necessary to probe whether the protospacer matches the guide RNA (PAM investigation state). If the protospacer is not correct, the formation of a partial R-loop is terminated and the dsDNA double helix is restored. The nuclease then reverts to the 1D state, in search of the next PAM motif. On the other hand, if the gRNA matches the sequence after the PAM, a complete R-loop is generated and the nuclease enters the stable, target-bound state. By applying a MC-DDA-based analysis to the diffusional histograms, we sought to determine the rates governing the transitioning between each of the four states.

### *Spy*dCas9 spends more time than *Lb*dCas12a on non-target interactions with DNA

We first analysed the diffusional histogram of *Spy*dCas9, *Sp*RYdCas9, *Lb*dCas12a and imp*Lb*dCas12a when provided with a scrambled guide with no complementary protospacer in the cell. Fitting with MC-DDA was performed by simulating the nucleases moving linearly through their 3D, 1D and PAM-investigating states and yielded kinetic rates associated with transitioning between these states (Figure 2A-B).

**Figure 2.**
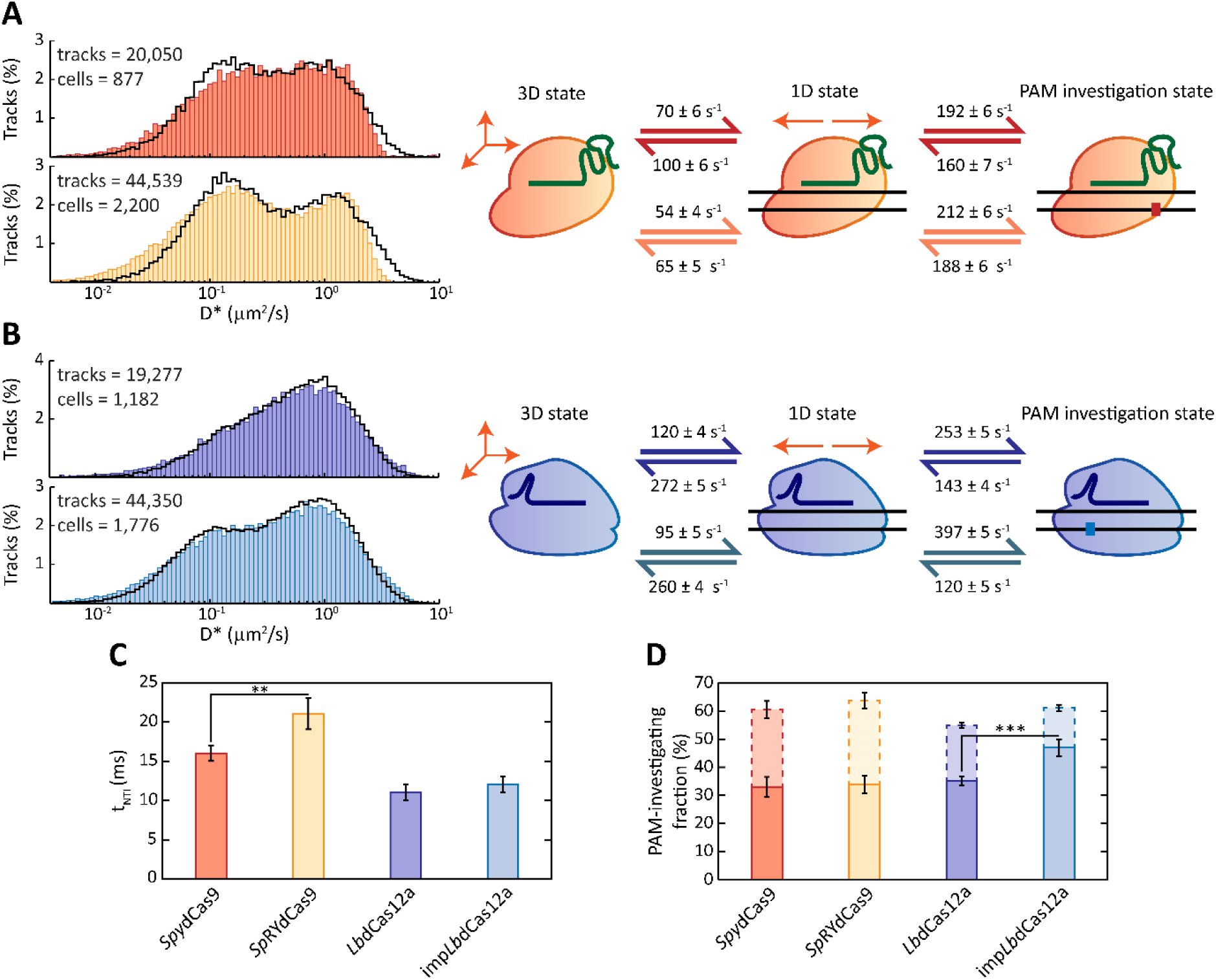
SptPALM of *Spy*dCas9, *Lb*dCas12a and their PAM-relaxed variant with a scrambled guide, without a complementary protospacer. **A)** On the left, diffusion coefficient histograms of tracks with a length of 3 steps (4 localisations) for *Spy*dCas9 (red, top) and *Sp*RYdCas9 (yellow, bottom). The number of tracks per histogram is shown as *n* in the upper left corner of each histogram. Histograms are fitted (black line) with a theoretical description of 250,000 particles moving between a 3D, a 1D and a PAM-investigating state. On the right, kinetic rates describing the fitting are provided for both *Spy*dCas9 (red arrows) and *Sp*RYdCas9 (orange arrows). **B)** On the left, diffusion coefficient histograms of tracks with a length of 3 steps for *Lb*dCas12a (purple, top) and imp*Lb*dCas12a (blue, bottom). The number of tracks per histogram is shown as *n* in the upper left corner of each histogram. Histograms are fitted as for panel A. On the right, kinetic rates describing the fitting are provided for both *Lb*dCas12a (purple arrows) and imp*Lb*dCas12a (blue arrows). **C)** Non-target interaction times (*t*_NTI_, in ms) for each of the four nucleases, calculated as 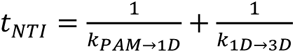. The two asterisks (**) indicate a statistically significant difference between the *Spy*dCas9 and the *Sp*RYdCas9 *t*_NTI_, with statistic performed by unpaired *t*-test, p < 0.01. Source data is available in Supplementary Material (Source Data Table 1). **D)** Fraction of the population of each of the four nucleases that is on average investigating a PAM sequence (bold bars) or in a 1D state (dashed bars), calculated as mentioned in the Supplementary Material (Initialisation of simulated particles). The three asterisks (***) indicate a statistically significant difference between the PAM-investigating fractions, with statistic performed by unpaired *t*-test, p < 0.001. All statistics in this figure were performed via unpaired *t-*test. Source data of panel is available in Supplementary Material (Source Data Table 2).

In our model, we assigned rates for the transition between the 1D and the other states (*k*_1D→3D_, *k*_1D→PAM_) to interpret the distribution obtained from sptPALM. However, while imaging, only the totality of the interaction of a dCas nuclease with the DNA can be confidently resolved and quantified. Therefore, the non-target DNA interaction time (*t*_NTI_) of each nuclease is expressed as a sum of the times needed to investigate a PAM and to fully detach from the DNA 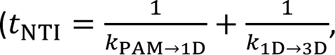 expressed in milliseconds) (Figure 2C). In this way, we obtain a t_NTI_ of 16 ± 1 ms for *Spy*dCas9. This value is in agreement with previous reports of an upper limit of 30 ms in *E. coli*^32^ and of ∼17 ms in either *L. lactis* cells^27^ or *in vitro* single-molecule experiments^21^. Interestingly, the PAM-relaxed variant *Sp*RYdCas9 exhibits a slightly increased retention on non-target DNA, with a *t*_NTI_ of 21 ± 2 ms. Several variants of the type V nuclease dCas12a were reported to exhibit faster interaction times in *in vitro* single-molecule settings than *Spy*dCas9^22^. Here, we show that this characteristic is maintained in live cells, with obtained rates describing overall fast dynamics between *Lb*dCas12a variants and non-target DNA. From these rates, we were able to obtain a *t*_NTI_ of 11 ± 1 ms for the wild-type nuclease and a value of 12 ± 1 ms for its imp*Lb*dCas12a variant.

The fraction of nucleases investigating PAM sequences on average (Figure 2D, bold bars) is roughly constant (∼33%) for both studied variants of *Spy*dCas9. On the other hand, relaxing the PAM requirements of the *Lb*dCas12a nuclease led to a significant increase in the PAM-investigating fraction from 35 ± 2% to 47 ± 3% in imp*Lb*dCas12a. In all cases, the total DNA-interacting fraction is close to the previously reported range of 55 - 89% for proteins with DNA target search properties in *E. coli*^56^. To further investigate the behaviour of the four nucleases while searching for a DNA target site, we generated diffusion coefficient histograms belonging to tracks with progressively increasing length and calculated the average diffusion coefficient (Figure 3)

**Figure 3.**
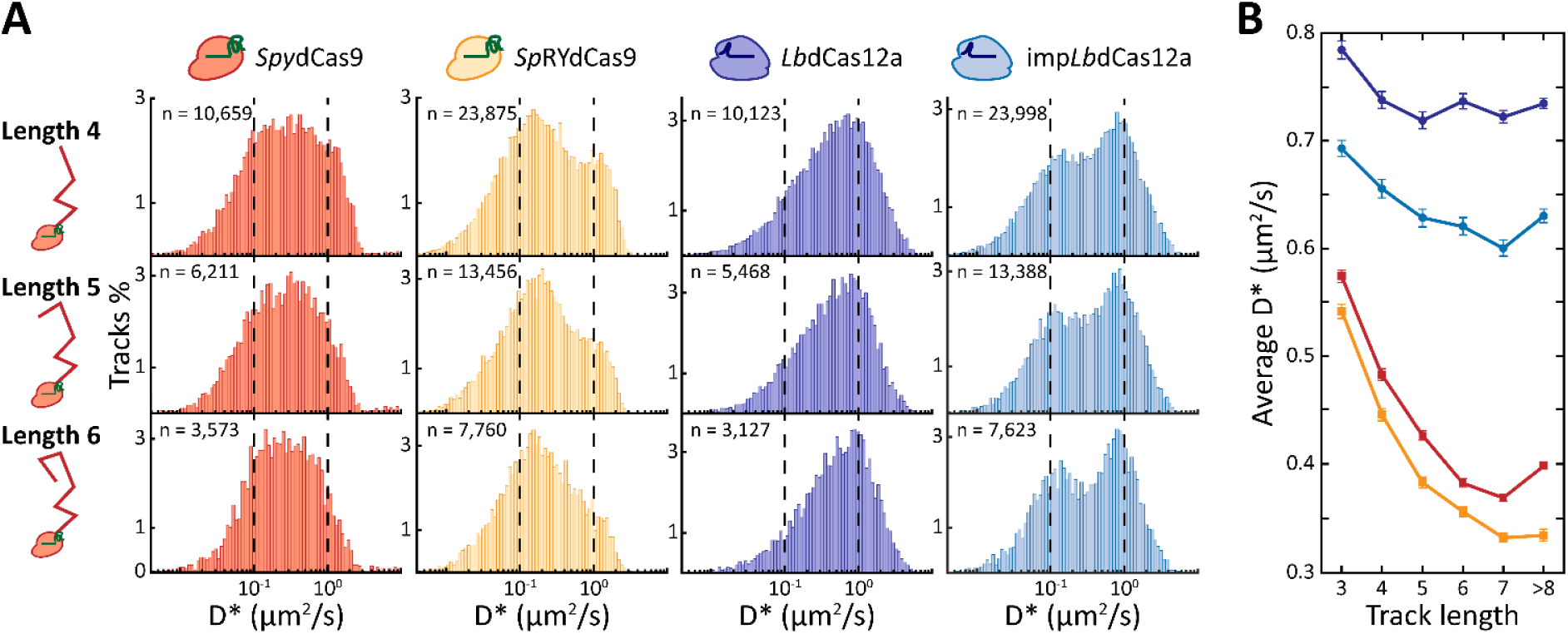
The behaviour of *Spy*dCas9, *Lb*dCas12a and their PAM-relaxed variants at increasing track length. **A)** Tracks belonging to the four nucleases with a length of either 4 steps (5 localisations), 5 steps (6 localisations) or 6 steps (7 localisations) are shown. The number of tracks per histogram is shown as *n* in the upper left corner of each histogram. The vertical dashed lines indicate diffusion coefficients of either 10^-1^ (left) or 10^0^ μm^2^/s (right) and are for visual aid. **B)** Trend of average diffusion coefficient at increasing track lengths (3 to ≥8) for *Spy*dCas9 (red line), *Sp*RYdCas9 (orange line), *Lb*dCas12a (purple line) and imp*Lb*dCas12a (blue line). Dots indicate the average *D*\* for track of the specific length for *Lb*dCas12a and imp*Lb*dCas12a, squares indicate the average *D*\* for *Spy*dCas9 and *Sp*RYdCas9, lines indicate the trend and the error bars indicate standard deviation of the histogram means. Source data of panel B is available in Supplementary Material (Source Data Table 3).

Overall, *Lb*dCas12a variants exhibit higher average diffusion coefficients than *Spy*dCas9 variants throughout all tested track lengths, highlighting once more their faster DNA probing in absence of a guide-matching protospacer. Interestingly, the average diffusion coefficient of both *Spy*dCas9 and *Sp*RYdCas9 considerably decreases for tracks with a higher number of localisations. For the wild-type *Spy*dCas9 nuclease, the average diffusion coefficient reduces from 0.57 ± 0.01 μm^2^/s for tracks with 3 steps (4 localisations) to 0.40 ± 0.00 μm^2^/s for tracks of more than 8 steps (≥9 localisations). The PAM-relaxed variant *Sp*RYdCas9 appears as less mobile on average, with averages ranging from 0.54 ± 0.01 μm^2^/s (track length 3) to 0.33±0.01 μm^2^/s (track length ≥8). On the other hand, *Lb*dCas12a is more mobile, with an average diffusion coefficient of 0.79 ± 0.01 μm^2^/s for tracks with a length of 3 steps. This value slightly decreases for tracks with an additional localisation, before remaining roughly constant around a value of ∼0.72 μm^2^/s. Strikingly, PAM-relaxed imp*Lb*dCas12a nuclease exhibits a mixed behaviour. The average diffusion coefficients obtained for this nuclease for different length tracks are overall higher than the ones of the studied *Spy*dCas9 nucleases, yet they follow a similar decreasing trend for longer tracks. Overall, our analysis revealed that, when compared to *Spy*dCas9, *Lb*dCas12a variants are considerably more dynamic in their way of probing the bacterial genome for matching protospacers.

### *Lb*Cas12a is predicted to find DNA target sites faster than *Spy*Cas9

After having obtained the kinetic rates governing the transition between the different states of our linear model, we set out to estimate what is the impact of the number of PAM sequences on the target search of each nuclease. To this end, we adapted a simulation used to predict *Spy*Cas9 cleavage in *L. lactis*^9^ for our linear four state model. We modified the model to accept different genomes of different sizes, copy number and GC content and different nucleases as input (Figure S3). At least 2,000 cells were simulated, each containing 1 or 10 units of either *Spy*dCas9, *Sp*RYdCas9, *Lb*dCas12a and imp*Lb*dCas12a. Each unit was initialised in a 3D state and given a set time interval to find a single target available inside the cell. The likelihood that the search was successful was obtained and then plotted as a function of the given search time (Figure 4A).

**Figure 4.**
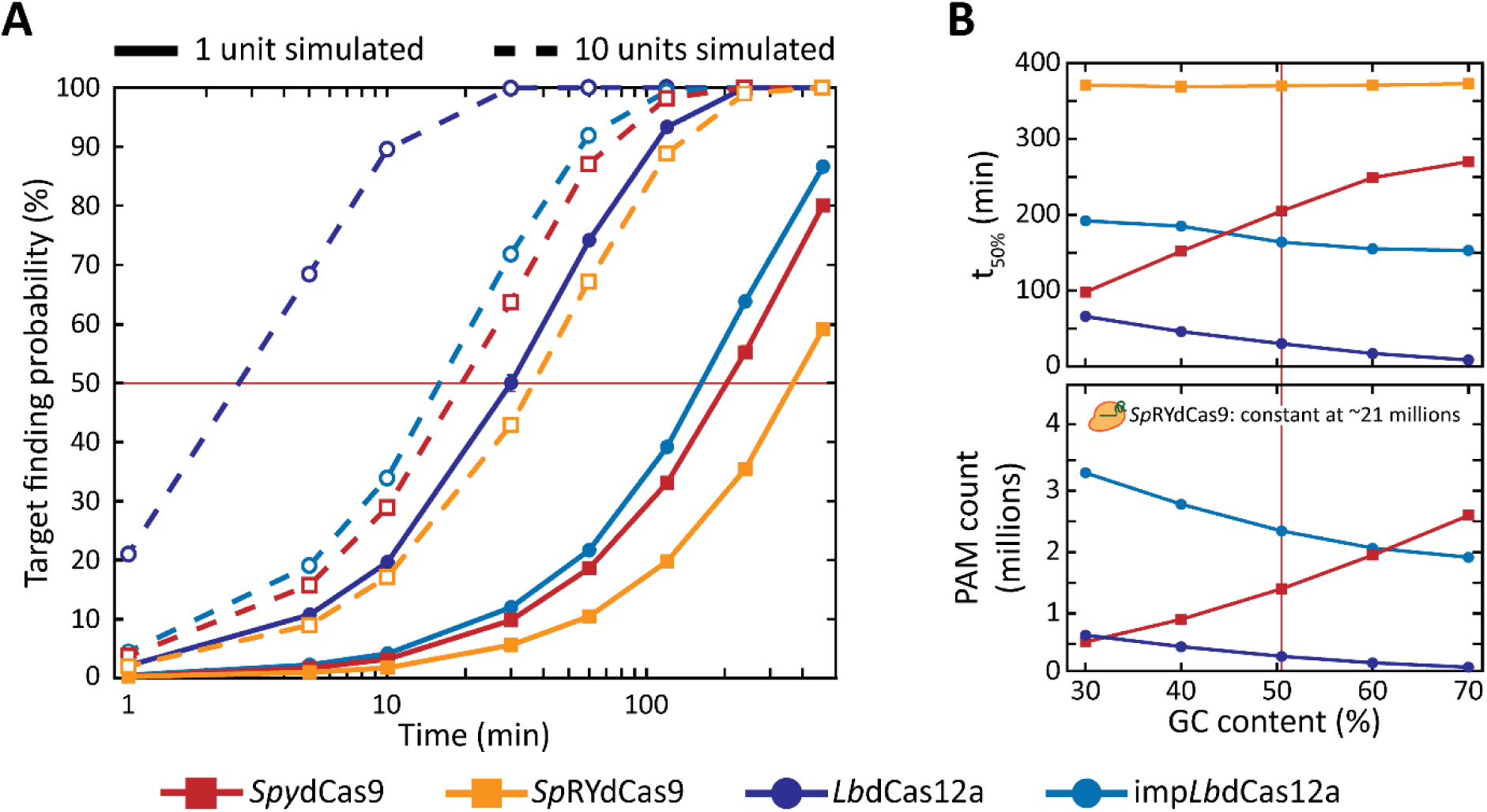
Simulation of DNA target search by *Spy*dCas9, *Sp*RYdCas9, *Lb*dCas12a and imp*Lb*dCas12a. **A)** Calculated probabilities of either one or ten units of *Spy*dCas9 (red line), *Sp*RYdCas9 (orange line), *Lb*dCas12a (purple line) or imp*Lb*dCas12a (blue line) finding and irreversibly binding to one target site in a *E. coli* cell after a certain time. Red line indicates 50% of target finding probability. Error bars indicate standard deviation calculated from ten iterations of the model. An explanation and list of the variables used is available in Supplementary Material (Variables used in the estimation of target search time of Cas nucleases). Source data is available in Supplementary Material (Source Data Table 4 and 5). **B)** Predicted time needed to find a target with 50% probability (*t*_50%_) (top) and corresponding number of PAM sequences (bottom) for a variety of GC contents. The size and copy number of the genome, together with size and copy number of the pTarget plasmid, were kept constant while varying GC content. Red line indicates the GC content of *E. coli* (50.5%). Source data is available in Supplementary Material (Source Data Table 6 and 7).

The simulation shows an inverse dependency between the number of PAMs and the target search time for almost all nucleases. A single *Spy*dCas9 is expected to have a 50% chance of finding a single target site after ∼200 min, whereas the PAM-relaxed *Sp*RYdCas9 takes almost double the time (∼370 min). In the same way, the predicted time of *Lb*dCas12a is only ∼30 min, against the ∼160 min needed for imp*Lb*dCas12a. Interestingly, imp*Lb*dCas12a is still faster than *Spy*dCas9 (∼160 min against ∼200 min), despite the engineered *Lb*dCas12a nuclease having almost one million more motifs to investigate. When the number of simulated proteins is increased by a factor of ten, the observed trends are conserved. The effect of multiple nucleases leads to a much higher reliability in finding a single target, as previously reported for *Spy*dCas9 in *L. lactis* cells^9^.

We further investigated how the number of PAM sequences could affect the activity of each protein by varying the GC content of the host while maintaining the other variables constant (i.e., size and copy number of genome, number of targets). We then obtained the time that *Spy*dCas9, *Lb*dCas12a and their PAM-relaxed variants take to find their target with a 50% probability (*t*_50%_) (Figure 4B). Due to its degenerated PAM motif, *Sp*RYdCas9 was not affected by the change in GC content and its *t*_50%_ remained constant. As expected, the increase in GC content had opposite effects on *Spy*dCas9 and *Lb*dCas12a, due to their preferences for G-rich and T-rich PAMs, respectively. *Spy*dCas9 ranged from a *t*_50%_ of ∼98 min to one of ∼270 min. On the other hand, *Lb*dCas12a needed ∼66 min to find its target with 50% of probability at 30% GC content, while this value dropped to ∼8.5 min when the GC content increased to 70%. Strikingly, the *t*_50%_ of the PAM-relaxed imp*Lb*dCas12a exhibited the smallest variation (aside for *Sp*RYdCas9), despite the number of available PAM sequences differing by almost 1.5 million among the extremes of the tested range. Our simulations thus revealed that the faster interactions of *Lb*dCas12a variants favour them in the recognition of a target sequence in most cellular scenarios.

### Relaxation of PAM requirements impacts target search of both *Spy*dCas9 and *Lb*dCas12a

Next, we performed sptPALM on cells with Cas nucleases harbouring a targeting guide. We replaced the scrambled, non-targeting guide on each pCas plasmid with a targeting guide, complementary to the five protospacers harboured on the pTarget plasmid. The heterologous *pLtetO-1* promoter used to drive expression of the different Cas nucleases resulted in heterogeneous protein expression in our cell populations. We therefore split the imaged cells into smaller sets according to the number of visualised tracks and investigated the effect that track numbers have when a finite number of protospacers is present (Figure 5A).

**Figure 5.**
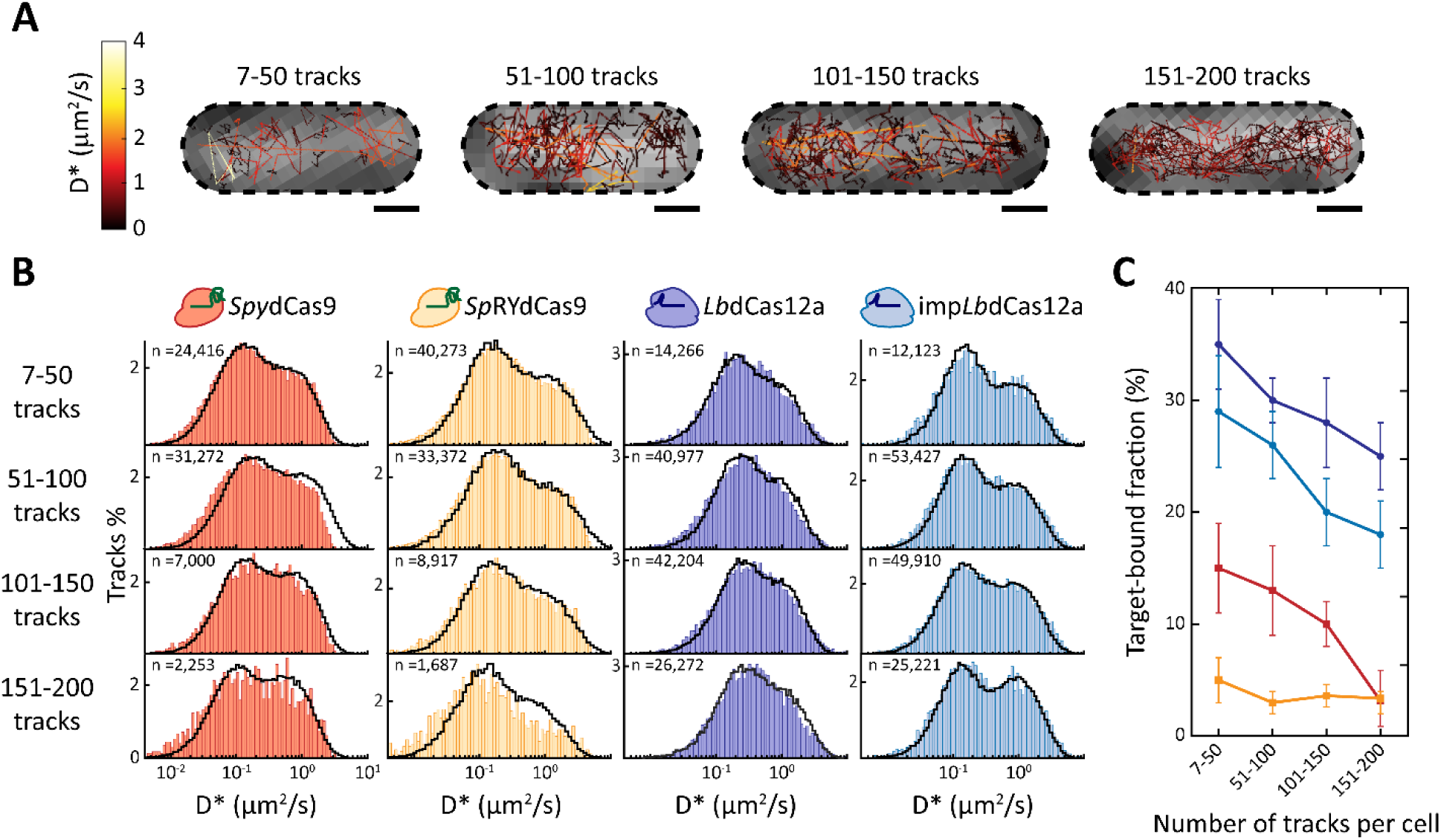
SptPALM of *Spy*dCas9, *Lb*dCas12a and their PAM-relaxed variant with a targeting guide, complementary to the protospacers on the pTarget plasmid. **A)** Visualised tracks in *E. coli* PAM-SCANR cells, harbouring one of a pCas-PAmCherry2.1 with a targeting guide and the pTarget plasmid. Tracks are coloured according to their apparent diffusion coefficient. Four cells are showed, belonging to the different ranges of copy numbers considered (7-50, 51-100, 101-150 and 150-200 tracks). Black, dashed lines indicate cell membranes, as determined from the brightfield images. Scale bar is 500 nm. **B)** Diffusion coefficient histograms of each nuclease for each range of track counts (85 bins). The number of tracks per histogram is shown as *n* in the upper left corner of each histogram. Black lines are the result of a fit performed with two species: a static species moving at the diffusion coefficient of the pTarget plasmid, corresponding to the target-bound nuclease; a three-state species, corresponding to the target-searching nucleases, with transition between states dictated by kinetic rates obtained for the scrambled distribution of the same nuclease. **C)** The target-bound fraction of each nuclease for each track count range, expressed as the percentage of the immobile species over the total number of particles simulated for the fit in panel B. Dots indicate the average target-bound fractions of *Lb*dCas12a and imp*Lb*dCas12a, squares indicate the average target-bound fractions of *Spy*dCas9 and *Sp*RYdCas9, lines indicate the trend and the error bars indicate standard deviation. Source data of panel C is available in Supplementary Material (Source Data Table 8).

In its target-bound state, a Cas nuclease is expected to move with a diffusion coefficient dictated by the size of the pTarget plasmid. We considered Cas nucleases interacting with their protospacers as an additional species moving with a diffusion coefficient dictated by the plasmid they are bound to. In agreement with diffusion coefficients previously estimated for plasmids^57,58^, this led to a 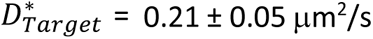. We thus fitted the obtained diffusion coefficients with a combination of this target-bound species and a target-searching species (Figure 5B). The latter moves in the linear three-state model previously described for nucleases carrying a scrambled guide and uses the kinetic rates obtained previously (Figure 2A,B). We obtained the fraction of Cas nucleases bound to their protospacers as a fraction of the total number of particles simulated (Figure 5C). In accordance with our previous work^9^, we observed that the fraction of *Spy*dCas9 molecules bound to their target site progressively decreased in cells with progressively higher numbers of visible tracks. This value was estimated at 15% for cells carrying 7 to 50 tracks and at only 3% when the track number increased to 151 to 200 tracks. All nucleases were found to exhibit a similar trend, with the sole exception of the PAM-relaxed *Sp*RYdCas9, for which the target-bound fraction stays roughly constant around 3%. Interestingly, relaxing PAM requirements led to a less efficient target search for both *Spy*dCas9 and *Lb*dCas12a. This is indicated by the generally lower bound fraction at every track number range.

## 4. Discussion

In this study, we designed and performed a sptPALM assay to compare the behaviour of *Spy*dCas9, *Lb*dCas12a and their PAM-relaxed variants (*Sp*RYdCas9 and imp*Lb*dCas12a) in live *E. coli* cells at the single-molecule level. We expanded our previous analysis of *Spy*dCas9 target search^9^ to include four states (3D state, 1D state, PAM-investigating state and target-bound state) rather than the previously considered three states. Because of this, we were able to extract kinetic rates governing the transition between the different states and to simulate the target search process of these genome editing tools in a variety of different backgrounds. We thus report not only the first single-molecule characterisation of *Lb*dCas12a, *Sp*RYdCas9 and imp*Lb*dCas12a in live cells, but also the first comparison at this level of some of the most employed genome editing tools.

### *Spy*dCas9 and *Lb*dCas12a have different rate limiting steps in PAM detection

When using a non-targeting guide, previous *in vitro* characterisation of *Spy*Cas9^11,21^ and *Lb*Cas12a^22^ DNA probing behaviour was in overall good agreement with our *in vivo* analysis. Notably, *Lb*dCas12a was observed to spend less time than *Spy*dCas9 on non-target interactions with DNA (11 ± 1 ms against 16 ± 1 ms, respectively). We attributed this difference to the reported ability of *Spy*dCas9 to slide between several adjacent PAM sequences before releasing from the DNA^11,21^. After encountering a random stretch of DNA, a *Spy*dCas9 molecule proceeds unidirectionally on DNA to find a PAM sequence. Due to the 50.5% GC content of the *E. coli* genome and the requirement of a 5’-NGG PAM, this sequence is encountered at almost every interaction with DNA. If the sequence immediately downstream of the PAM is not complementary to the gRNA, the nuclease has a probability to move to nearby PAM motifs^11,21^. The likelihood of transitioning to adjacent PAMs is mostly dependent on the distance between them. The number of sequences investigated sequentially then dictates the total amount of time spent on the DNA, here expressed by the *t*_NTI_ value. This finding explains the increase of the obtained *t*_NTI_ from the 16 ± 1 ms of *Spy*dCas9 to the 21 ± 2 ms of the PAM-relaxed *Sp*RYdCas9, which now recognises every stretch of three nucleotides as a PAM sequence. Following this model of action, we can assume that the rate-limiting step in investigating one or more PAM sequences for both variants of *Spy*dCas9 is the first, non-specific interaction with the DNA. This assumption is supported by two other experimental observations. First, a decrease in average diffusion coefficients for progressively increasing track lengths was observed for both *Spy*dCas9 variants (Figure 3B). The decrease suggests that the longer a single nuclease is observed, the more likely it is to be found in a DNA-bound state. This observation could be explained by the increased impact of blurred PSFs at longer track lengths^59^. Yet, the consistently high mobility of *Lb*dCas12a suggests that the analysis is not excessively biased by motion blur. This finding indicates relatively long-lived interactions with DNA after the first random transition from the 3D to the 1D state. Second, the comparable PAM-investigating fractions of *Spy*dCas9 and *Sp*RYdCas9 (Figure 2D) further suggests that at least one sequence is always investigated for a match with the spacer at every contact with DNA.

*Lb*dCas12a showed a different behaviour when equipped with a scrambled guide. First, we observed no difference in the obtained *t*_NTI_ between the wild-type *Lb*dCas12a (11 ± 1 ms) and the PAM-relaxed variant (12 ± 1 ms). Moreover, the average diffusion coefficient of the *Lb*dCas12a remained overall constant across different track lengths, whereas the one of imp*Lb*dCas12a only slightly decreased (from 0.69 μm^2^/s for track length 3 to a minimum of 0.60 μm^2^/s for track length 7). This data can point to the lack of shuttling between PAMs for *Lb*dCas12a, since there is no striking reduction in mobility even when the expected number of PAM sequences is increased almost 10-fold (from 2.4⋅10^5^ to 2.3⋅10^6^ in our genetic background). The slight decrease in mobility of imp*Lb*dCas12a could also be attributed to off-targets in the presence of a scrambled guide. Our MC-DDA analysis does not take off-target events into account and imp*Lb*Cas12a has already been deduced to result in more off-target events than to its wild-type counterpart from editing frequencies analysis^25^. The relaxation of the PAM requirements, on the other hand, led to a significant increase from 35% to 47% in the fraction of *Lb*dCas12a nucleases investigating PAM sequences (Figure 2D). Taken together, these results suggest that the limiting factor in investigating a sequence for a correct spacer match is not random contact with DNA as with *Spy*dCas9, but rather encountering a correct PAM sequence. The overall higher average diffusion coefficient for both *Lb*dCas12a variants in our *in vivo* analysis also indicates shorter interactions with the DNA and might suggest a confirmation of a previous reported hopping motion^12,28^.

### Both kinetic rates and number of PAM sequences impact target search

The kinetic rates obtained through MC-DDA analysis of our diffusional histogram with a scrambled guide allowed us to simulate the target search process. We simulated either 1 or 10 units of each Cas nuclease searching for a single target in the same genetic background of our experimental setting (Figure 4A). In this way, we predicted *Lb*dCas12a to be the fastest nuclease, with a t_50%_ value of ∼30 min. This means that a single protein of this nuclease has a 50% probability to find and potentially cleave a single target sequence in roughly the same average time it takes an *E. coli* cell to divide^49^. As a comparison, the LacI repressor was experimentally determined to need 5 min to find its operator sequence in *E. coli*^60,61^. However, it is important to notice that the Cas nuclease target search process resembles the transcription factor one only up to the PAM investigation state. The rate-limiting step is finding a protospacer among all the wrong sequences that are preceded by a PAM. The reported speed of *Lb*dCas12a is surely helped by the favourable GC content of this bacterium (50.5%). In this context, there is a relatively low number of PAM sequences to investigate for a nuclease that prefers T-rich motifs. The impact that the number of PAMs has on the target search process is particularly evident for the PAM-relaxed *Sp*RYdCas9, where the *t*_50%_ is the highest of all the tested nucleases at ∼370 min. However, our simulation led to the conclusion that the number of available PAM sequences is not the sole determinant of the target search efficiency of a nuclease. Despite the higher number of recognised motifs, the overall faster kinetics of imp*Lb*dCas12a led to a faster target search (∼164 min, with 2.3⋅10^6^ PAM motifs) compared to the wild-type *Spy*dCas9 (∼205 min, with 2.4⋅10^5^ PAM motifs). More interestingly, when simulated across a range of different GC contents and thus number of PAMs, *Spy*dCas9 was predicted to be faster than imp*Lb*dCas12a only at the lower GC contents (30% and 40%). In these conditions, however, the number of available sequences to investigate for imp*Lb*dCas12a was always at least 2 million higher than for *Spy*dCas9. This result suggests that faster target search kinetics are almost always favourable in identifying one correct sequence from a vast majority of non-target ones. It is thus interesting to ask why throughout evolution both modes of action were established in the different Cas nucleases. The *Lb*dCas12a ancestor protein TnpB has already been characterised^62^ and reprogrammed for genome editing purposes^63^, similarly to the *Spy*dCas9 ancestor IscB^64–66^. The emergence of specific target search modes could thus be investigated by applying our sptPALM assay to both TnpB and IscB.

Our assessment of target-bound fraction of Cas nucleases harbouring a targeting guide (Figure 5) showed a similar trend to the predicted target search times. When fitting diffusional histograms derived from cells containing 7-50 tracks, *Lb*dCas12a had the highest observed target-bound fraction (30%). The wild-type *Lb*dCas12a was followed by its PAM-relaxed variant (29%) and *Spy*dCas9 (15%). Finally, only 5% of imaged *Sp*RYdCas9 was bound to its target in the presence of a protospacer. This low value is likely a combination of both the massive amount of sequences to investigate and the tendency of *Sp*RYdCas9 to cluster on non-target DNA loci, as recently reported^67^. Once again, it is striking to notice how imp*Lb*dCas12a is more efficient in finding its target than wild-type *Spy*dCas9, despite the Cas12a variant having a higher number of PAM sequences and a lowered specificity^25^. The general difference between the values obtained for each nuclease could have several reasons.

Notably, replication forks can remove proteins from the DNA, including *Spy*dCas9^32^. If we assume that the full genome of *E. coli*, including the pTarget plasmid, is replicated at intervals of the doubling time, then in our experimental settings all proteins are moved to their 3D state by DNA replication with an interval of 40 min as lower bound. The removal of Cas proteins from DNA does not only happens at regular intervals through replication, but is also mediated in *E. coli* more regularly by cellular factors such as the RNA polymerase and Mdf translocase^68^. A faster target search process might then lead to an overall higher bound fraction in the steady-state conditions of our turbidostat-cultivated cells. In all cases, the target-bound fraction decreased when the number of imaged proteins per cell increased, suggesting competition for a limited number of available targets. Moreover, all the theoretically available protospacers (5 protospacers/plasmid ⋅ ∼64 plasmids^40^ = ∼320 protospacers) were not filled for any nuclease nor track per cell range. The lack of occupancy of all available targets can be explained by a decrease in the overall number of plasmids due to Cas interference^9^ or steric hindrance. DNA footprints of *Spy*dCas9 and a different Cas12a variant (*Fn*dCas12a) were reported to be ∼78 bp^69^ and ∼35 bp^70^, respectively. Each target site on our pTarget plasmid is distanced by the adjacent one by 30 bp. Therefore, both Cas proteins could mask key features of a protospacer in the vicinity of the one it is currently bound to. The steric hindrance, in turn, reduces the overall number of available sites and thus our target-bound fraction.

Overall, the here-reported *in vivo* data supports previous *ex situ* characterisation of *Spy*dCas9^11,21^ and *Lb*dCas12a^12,21,28^ and expands the knowledge on target search in live cells. *Spy*dCas9 is once again reported to spend more time on non-target interaction with DNA than *Lb*dCas12a^22,28^. As a consequence, the latter nuclease finds its target faster in our simulations. The relaxation of PAM requirements generally leads to a less efficient target search for both nucleases, as deduced from the fraction of proteins bound to their target at any given moment and predicted target finding times. Despite the impact on target search efficiency, the faster dynamics of imp*Lb*dCas12a still led to a higher, observed target-bound fraction in our sptPALM assays. The target search simulations presented here might also suggest how the same nucleases would behave in different organisms. Altogether, we characterised rate-limiting steps in PAM recognition for both *Spy*Cas9 and *Lb*Cas12 nucleases and their variants. We then demonstrated how the increase in flexibility due to the relaxation of PAM requirements negatively impacts the efficiency of target search in live cells for both *Spy*dCas9 and *Lb*dCas12a. The framework presented here provides a powerful basis for future characterisation of established or new Cas nuclease variants or other DNA targeting proteins.

## Data availability

All data used in this study to generate plots in the various figure is available as Source Data in a corresponding table, as mentioned in the caption of each figure. All the localisation data, as well as the scripts used in the analysis are available on Zenodo (https://doi.org/10.5281/zenodo.10142521). All other data, together with all the plasmid sequences are available upon request by contact with the corresponding authors.

## Acknowledgements

L.O. and J.vd.O. acknowledge financial support from The Netherlands Organization of Scientific Research (NWO/OCW) Gravitation program Building A Synthetic Cell (BaSyC) (024.003.019). R.H.J.S. is supported by a VIDI grant (VI.Vidi.203.074) from NWO.

The authors would like to thank Mink Neeleman, Tim Hendriksen and Thijs Hermans for their help in the early stages of the project.

