## Supplementary material, Supplementary information and Source Data for "Live-cell imaging reveals the trade-off between target search flexibility and efficiency for Cas9 and Cas12a"

**Table S1. List of bacterial strains used in this study.**

| Strains | Characteristics | Source |
|---|---|---|
| *E. coli* DH5a | $F^-$ *endA1 glnV44 thi-1 recA1 relA1 gyrA96 deoR nupG purB20* φ80d*lacZ*ΔM15 Δ(*lacZYA-argF)*U169 *hsd*R17(*$r_K$– $m_K$+*), l$^-$. | NEB |
| *E. coli* PAM-SCANR | *lacIq* *$rrnB_{T14}$* Δ*$lacZ_{WJ16}$* *$hsd_{R514}$* Δ*$araBAD_{AH33}$* Δ*$rhaBAD_{LD78}$* *rph-1* Δ*cas3*-CRISPR1 Δ$P_{lacI}$-*lacZ*. | [32] |
| *E. coli* PAM-SCANR x p*Spy*dCas9-PAmCherry2.1_Target, pTarget | PAM-SCANR strain harbouring plasmids p*Spy*dCas9-PAmCherry2.1_Target and pTarget. | This study |
| *E. coli* PAM-SCANR x p*Spy*dCas9-PAmCherry2.1_Scrambled, pTarget | PAM-SCANR strain harbouring plasmids p*Spy*dCas9-PAmCherry2.1_Scrambled and pTarget. | This study |
| *E. coli* PAM-SCANR x p*Lb*dCas12a-PAmCherry2.1_Target, pTarget | PAM-SCANR strain harbouring plasmids p*Lb*dCas12a-PAmCherry2.1_Target and pTarget. | This study |
| *E. coli* PAM-SCANR x p*Lb*dCas12a-PAmCherry2.1_Scrambled, pTarget | PAM-SCANR strain harbouring plasmids p*Lb*dCas12a-PAmCherry2.1_Scrambled and pTarget. | This study |
| *E. coli* PAM-SCANR x p*Sp*RYdCas9-PAmCherry2.1_Target, pTarget | PAM-SCANR strain harbouring plasmids p*Sp*RYdCas9-PAmCherry2.1_Target and pTarget. | This study |
| *E. coli* PAM-SCANR x p*Sp*RYdCas9-PAmCherry2.1_Scrambled, pTarget | PAM-SCANR strain harbouring plasmids p*Sp*RYdCas9-PAmCherry2.1_Scrambled and pTarget. | This study |
| *E. coli* PAM-SCANR x pimp*Lb*dCas12a-PAmCherry2.1_Target, pTarget | PAM-SCANR strain harbouring plasmids pimp*Lb*dCas12a-PAmCherry2.1_Target and pTarget. | This study |
| *E. coli* PAM-SCANR x pimp*Lb*dCas12a-PAmCherry2.1_Scrambled, pTarget | PAM-SCANR strain harbouring plasmids pimp*Lb*dCas12a-PAmCherry2.1_Scrambled and pTarget. | This study |

**Table S2. List of plasmids used in this study.**

| Plasmid | Relevant characteristics | Source |
|---|---|---|
| pBeloBAC11 | *repE, ori2, sopABC, cat.* | NEB |
| pSC101-K102E | *neoR, rep101(K102E).* | [37] |
| pd*Spy*Cas9-PAmCherry2.1_Target | pBeloBAC11, $P_{LtetO-1}$-BCD-*Spydcas9-PAmCherry2(M10L)*, $P_{J23119}$-*sgRNA(targeting)*. | This study |
| pd*Spy*Cas9-PAmCherry2.1_Scrambled | pBeloBAC11, $P_{LtetO-1}$-BCD-*Spydcas9-PAmCherry2(M10L)*, $P_{J23119}$-*sgRNA(scrambled)*. | This study |
| pd*Lb*Cas12a-PAmCherry2.1_Target | pBeloBAC11, $P_{LtetO-1}$-BCD-*Lbdcas12a-PAmCherry2(M10L)*, $P_{J23119}$-*sgRNA(targeting)*. | This study |

| | | |
|---|---|---|
| pd*Lb*Cas12a-PAmCherry2.1_Scrambled | pBeloBAC11, $P_{LtetO\text{-}1}$-BCD-*Lbdcas12a--PAmCherry2(M10L)*, $P_{J23119}$-*sgRNA(scrambled)*. | This study |
| pSpRYdCas9-PAmCherry2_Target | pBeloBAC11, $P_{LtetO\text{-}1}$-BCD-*Spydcas9(A61R, L1111R, D1135L, S1136W, G1218K, E1219Q, A1322R, R1333P, R1335Q, T1337R)-PAmCherry2(M10L)*, $P_{J23119}$-*sgRNA(targeting)*. | This study |
| pSpRYdCas9-PAmCherry2_Scrambled | pBeloBAC11, $P_{LtetO\text{-}1}$-BCD-*Spydcas9(A61R, L1111R, D1135L, S1136W, G1218K, E1219Q, A1322R, R1333P, R1335Q, T1337R)-PAmCherry2(M10L)*, $P_{J23119}$-*sgRNA(scrambled)*. | This study |
| pimp*Lbd*Cas12a-PAmCherry2_Target | pBeloBAC11, $P_{LtetO\text{-}1}$-BCD-*Lbdcas12a(D156R, G532R, K538V, Y542R, K595R)-PAmCherry2(M10L)*, $P_{J23119}$-*sgRNA(targeting)*. | This study |
| pimp*Lbd*Cas12a-PAmCherry2_Scrambled | pBeloBAC11, $P_{LtetO\text{-}1}$-BCD-*Lbdcas12a(D156R, G532R, K538V, Y542R, K595R)-PAmCherry2(M10L)*, $P_{J23119}$-*sgRNA(scrambled)*. | This study |
| pTarget | pSC101-K102E, 5 protospacers, 5 30 bp-long distancing sequences. | This study |

**Table S3. List of oligonucleotides and DNA fragments used to introduce mutations.** Nucleotides in bold introduced the listed mutations.

| Identifier | Sequence (5'-3') | Used for |
|---|---|---|
| BG28281 | **AAG**gttatcctcctcgcccttgctc | Introducing M10L mutation in PAmCherry2.1. |
| BG28278 | agggcgaggaggataac**CTT**gccatcatcaaggagttcatgcg | Introducing M10L mutation in PAmCherry2.1. |
| BG30037 | cggaggcttctccaaggaaagtatc**CGC**ccgaaaaggaacagcgacaagctgatcgcacgcaaaaaagattgggaccccaagaaatacggcggattc**CTGTGG**cctacagtcgcttacagtgtactggttgtggccaaagtggagaaagggaagtctaaaaaactcaaaagcgtcaaggaactgctgggcatcacaatcatggagcgatcaagcttcgaaaaaaaccccatcgactttctcgaggcgaaaggatataaagaggtcaaaaaagacctcatcattaagcttcccaagtactctctctttgagcttgaaaacggccggaaacgaatgctcgctagtgcg**AAACAG**ctgcagaaaggtaacgagctggcactgccctctaaatacgttaatttcttgtatctggccagccactatgaaaagctcaaagggtctcccgaagataatgagcagaagcagctgttcgtggaacaacacaaacactaccttgatgagatcatcgagcaaataagcgaattctccaaaagagtgatcctcgccgacgctaacctcgataaggtgctttctgcttacaataagcacagggataagcccatcagggagcaggcagaaaacattatccacttgtttactctgacc**CGC**ttgggcgcgcct**CGC**gccttcaagtacttcgacaccaccatagac**CCG**aag**CAA**tac**CGT**tctacaaaggaggtcctggacg | Introducing L1111R, D1135L, S1136W, G1218K, E1219Q, N1317R, A1322R, R1333P, R1335Q and T1337R mutations of *Sp*RYdCas9. |

| | | |
|---|---|---|
| BG30041 | gt**GCG**ttcggccgtctccccg | Introducing A61R mutation of *Sp*RYdCas9. |
| BG30042 | ggggagacggccgaa**CGC**acgcggctcaaaagaacag | Introducing A61R mutation of *Sp*RYdCas9. |
| BG29090 | tatatatgatgcaattcgtaattacgtaacccaaaagccgtacagcaaagataagttcaaactgtatttccagaacccgcagtttatgCGCggctgggacaaagac**GTT**gagacagac**CGC**cgcgccactattctgcgttacggcagcaagtactatctcgccatcatggacaaaaaatatgcaaagtgtctgcagaaaatcgataaagacgacgtgaacggaaattacgaaaagattaattataagctgctgccagggcccaacaagatgttaccgCGCgtattttttccaaaaa | Introducing G532R, K538V, Y542R and K595R mutation of imp*Lb*dCas12a. |
| BG29087 | aacatattttccctgtt**GCG**gaaaaagcccgtgaaggc | Introducing D156R mutation of imp*Lb*dCas12a. |
| BG29088 | **CGC**aacagggaaaatatgttttcagagg | Introducing D156R mutation of imp*Lb*dCas12a. |

### Supplemental information

#### Fitting of the histogram derived from Cas nucleases with a scrambled guide

##### Determination of apparent diffusion coefficient of 3D, 1D and PAM-investigating states

During MC-DDA, initial diffusion coefficients and kinetic rates are provided to initiate a simulation. Several rounds of optimisation are then performed to obtain a fit of the experimental histogram, yielding rates governing the transition between each of the three states present when a Cas nuclease is provided with a scrambled guide. Table S4 lists the apparent diffusion coefficients D* of each state.

The diffusion coefficient of the nucleases moving in 3 dimensions ($D_{3D}$) was estimated based off their hydrodynamic radius. AlphaFold[1]/ColabFold[2] was used to predict the three-dimensional structures of the fusion proteins *Spy*dCas9-PAmCherry2.1 and *Lb*dCas12a-PAmCherry2.1 (Figure S1). The crystal structures obtained in this way were then used as input for the *in silico* tool HullRad[3]. This yielded an hydrodynamic radius of 5.11 nm for *Spy*dCas9-PAmCherry2.1 (corresponding to a diffusion coefficient of ~42 $\mu m^2/s$) and 5 nm for *Lb*dCas12a-PAmCherry2.1 (corresponding to a diffusion coefficient of ~44 $\mu m^2/s$). From here, cytoplasmic retardation was applied to obtain apparent diffusion coefficients $D^*_{3D}$ of 2 $\mu m^2/s$ and 2.2 $\mu m^2/s$ for *Spy*dCas9 and *Lb*dCas12a, respectively.

Regarding one-dimensional diffusion coefficients, we used previous reports on the dynamics of both nucleases obtained through *in vitro* single-molecule approaches. *Spy*dCas9 has been reported to unidirectionally search for a PAM sequence on the DNA by a sliding mechanism of an average length of 20 bp[4,5]. We calculated that the covered distance in this state is ~0.00068 $\mu m$ (considering an average of 0.000034 $\mu m$/bp[6]). This value is lower than our localisation error $\sigma$ (0.033 $\mu m$) and thus a single *Spy*dCas9 is apparently immobile while in its 1D sliding state ($D^*_{1D}$ = 0 $\mu m^2/s$). On the other hand, *Lb*dCas12a has been reported to search for a PAM through a hopping mechanism with an estimated D of 1.75 $\mu m^2/s$ [7]. Hopping motion consists of intermittent contact with the DNA and as such it can be already potentially described by our model through fast kinetic rates governing transitions between the 3D and the 1D state. Therefore, we also considered *Lb*dCas12a as apparently immobile while in its 1D sliding state ($D^*_{1D}$ = 0 $\mu m^2/s$).

Finally, after encountering a PAM, both *Spy*dCas9 and *Lb*dCas12a molecule stop on the DNA to investigate the protospacer sequence for a match with their gRNA. In this state, the two nucleases are thus immobile and their $D^*_{PAM}$ is 0 $\mu m^2/s$.

**Table S4 Diffusion coefficients and apparent diffusion coefficients (after cytoplasmic retardation) for *Spy*dCas9, *Lb*dCas12a and their PAM-relaxed variants.**

| Nuclease | State | D ($\mu m^2/s$) | Obtained from | D* ($\mu m^2/s$) |
|---|---|---|---|---|
| *Spy*dCas9, *Sp*RYdCas9 | 3D | 41.9 | Hydrodynamic radius | 2 |
| | 1D | 0 | Covered space | 0 |
| | PAM | 0 | Static interaction | 0 |
| *Lb*dCas12a, imp*Lb*dCas12a | 3D | 44.1 | Hydrodynamic radius | 2.2 |
| | 1D | 0 | Covered space | 0 |
| | PAM | 0 | Static interaction | 0 |

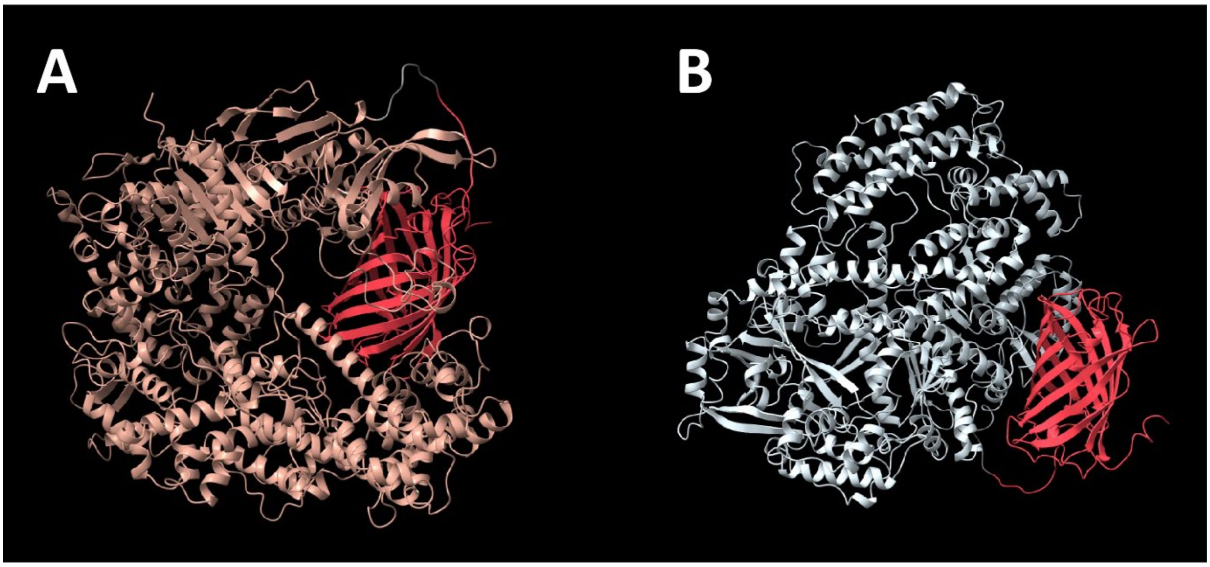


**Figure S1. AlphaFold-predicted 3D structures of A) *Spy*dCas9-PAmCherry2.1 and B) *Lb*dCas12a-PAmCherry2.1.** *Spy*dCas9 is coloured in light red, *Lb*dCas12a is coloured in light blue, the linker sequences are coloured in grey and the photoactivatable fluorescent protein PAmCherry2.1 is coloured in bright red.

#### Initialisation of simulated particles

At the beginning of MC-DDA, an initial set of kinetic rates ($k_{3D\to 1D}$, $k_{1D\to 3D}$, $k_{1D\to PAM}$, $k_{PAM\to 1D}$) are provided as an initial guess to interpret the distribution. These rates are used to determine the starting state of each simulated protein as follows:

$$c_{3D} = \frac{k_{PAM\to 1D} \cdot k_{1D\to 3D}}{(k_{1D\to PAM} \cdot k_{3D\to 1D}) + [k_{PAM\to 1D} \cdot (k_{1D\to 3D} + k_{3D\to 1D})]}$$

$$c_{1D} = \frac{k_{PAM\to 1D} \cdot k_{3D\to 1D}}{(k_{1D\to PAM} \cdot k_{3D\to 1D}) + [k_{PAM\to 1D} \cdot (k_{1D\to 3D} + k_{3D\to 1D})]}$$

$$c_{PAM} = \frac{k_{1D\to PAM} \cdot k_{3D\to 1D}}{(k_{1D\to PAM} \cdot k_{3D\to 1D}) + [k_{PAM\to 1D} \cdot (k_{1D\to 3D} + k_{3D\to 1D})]}$$

Where c is the probability of a protein being in the specific state, with $c_{3D} + c_{1D} + c_{PAM} = 1$.

#### Prediction of target search time

##### Variables used in the prediction of target search time of Cas nucleases

To estimate the probability that one of the studied Cas nucleases is able to find a single target in a specific cell, our software takes into consideration several variables. The user provides the time available to the nuclease, the number of units to simulate per cell, the GC content of the organism, the specific PAM sequence, the number of times to repeat the simulation, the size and average copy number of the genome, the size and copy number of the pTarget plasmid and the dissociation probability $P_{dis}$. This latter value is defined as $P_{dis} = \frac{1}{s^2}$, where s is the sliding length of the Cas nuclease in 1D motion. To estimate $P_{dis}$, we used the previously reported probabilities of *Spy*dCas9 transitioning between adjacent PAMs when the distance between these motifs was increased[4]. We fitted this data using exponential decay and obtained $P_{dis} \approx 0.144$ (Figure S2), corresponding to an average $s$ = 3 bp. Due to the lack of experimental characterisation of 1D sliding length for the remainder of the investigated proteins, we considered all Cas nucleases having the same average sliding length $s$ and $P_{dis}$ of *Spy*dCas9.

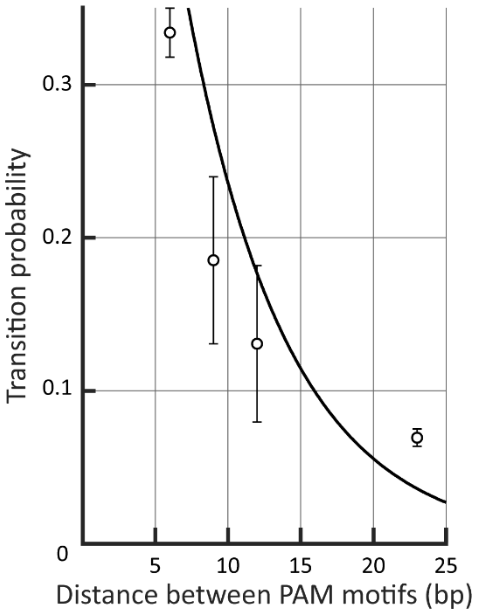


**Figure S2 Exponential decay fit of *Spy*dCas9 transition probability between adjacent PAM motifs.** The transition probabilities between PAM motifs with varying distances previously described[4] were fitted with an exponential decay to determine the $P_{dis}$ value (black line $y = e^{-0.144x}$).

From the input data, several other nuclease- and host-specific values are calculated and used (the complete set is available in Table S5). Two values are obtained through the GC content of the organism, the size of genome and pTarget plasmid and their average copy numbers: 1) the average number of PAMs; 2) the chance that a random stretch of N nucleotides matches the PAM sequence ($P_{PAM}$), where N is the PAM sequence length. Now knowing $P_{PAM}$, one can estimate the average number of PAMs a nuclease encounters when sliding on the DNA ($n_{1D\_PAM}$) for a length *s* (and thus a dissociation probability $P_{dis}$), through:

$$n_{1D\_PAM} = 1 + 4 \cdot P_{PAM} \cdot \sqrt{\sum e^{\ln(1-P_{dis})}}$$

Here, the sum corresponds to the average number of times the protein will move to the base situated left or right from the base it is currently on. Since it is a diffusional process, the square root is used to find the average sliding length, corresponding to the region where the protein searches for PAM sequences. One factor 2 is added to account for the effect that the protein can slide either way, and another factor 2 because the protein can bind on either one of two strands of the DNA.

**Table S5. List of values used for *Spy*dCas9, *Sp*RYdCas9, *Lb*dCas12a and imp*Lb*dCas12a for the simulation of target search.** For imp*Lb*dCas12a, recognising PAMs not collapsible in one notation (5'-TNTN, 5'-TACV, 5'-TTCV, 5'-CTCV, 5'-CCCV), the average number of each motif was obtained and the sum was used during the simulation. In the same way, the $P_{PAM}$ value used is a sum of the $P_{PAM}$ value of each individual PAM sequence.

| | **Cas nuclease** | | | |
|---|---|---|---|---|
| **Variables** | ***Spy*dCas9** | ***Sp*RYdCas9** | ***Lb*dCas12a** | **imp*Lb*dCas12a** |
| Genome size (bp) | 4,654,359 | 4,654,359 | 4,654,359 | 4,654,359 |
| Genome copies | 2.3 | 2.3 | 2.3 | 2.3 |
| pTarget size (bp) | 3,868 | 3,868 | 3,868 | 3,868 |
| pTarget copies | 1 | 1 | 1 | 1 |
| $P_{dis}$ | 0.144 | 0.144 | 0.144 | 0.144 |
| PAM sequence (5'-3') | 5'-NGG | 5'-NNN | 5'-TTTV | 5'-TNTN, 5'-TACV, 5'-TTCV, 5'-CTCV, 5'-CCCV |
| Average PAM count | 1,365,518 | 21,417,790 | 244,347 | 2,324,340 |
| $P_{PAM}$ | 0.0638 | 1 | 0.0114 | 0.1085 |
| $n_{1D_{PAM}}$ | 1.6201 | 10.7253 | 1.111 | 2.0552 |

#### Simulation of target search

Each nuclease unit was individually simulated as mentioned for the MCDDA of diffusional histograms of Cas proteins with scrambled guides. As mentioned before, a $t_{Change}$ times (Materials and methods: Monte-Carlo diffusion distribution analysis of Cas nucleases with a scrambled guide) and initial state of the simulated unit (Supplementary information: Initialisation of simulated particles) were assigned according to the specific rates for transitions between states. When in 1D state, the nuclease had three potential fates. A random number *p* (with 0 < *p* < 1) was generated and used to determine the fate of the unit.

First, a p ≤ $P_{target} = \frac{nr\ target\ sites - nr\ bound\ proteins}{nr\ PAM\ motifs} \cdot n_{1D_{PAM}}$ meant the nuclease found a PAM immediately followed by a matching protospacer, entering an irreversibly bound state. We only simulated the presence of 1 target site, thus ending the simulation for the current cell.

Second, if p ≤ $(c_{PAM} \cdot n_{1D_{PAM}})$ a PAM was encountered, yet not followed by the correct protospacer. A new $t_{Change}$ was then assigned according to the $k_{PAM \to 1D}$ rate and the simulation continued.

If none of the previous cases was true, the nuclease was considered as reverting to its 3D state after not having found any PAM sequence. Its $t_{Change}$ was thus changed according to the $k_{3D \to 1D}$ rate and the simulation continued.

### Source data

**Source Data Table 1. Source data of non-specific interaction times ($t_{NSI}$), used for the bar plot of Figure 3C.**

| Nuclease | $t_{NSI}$ (ms) | Standard deviation |
|---|---|---|
| *Spy*dCas9 | 16.25 | 0.659369446 |
| *Sp*RYdCas9 | 20.70376 | 1.195545762 |
| *Lb*dCas12a | 10.66948 | 0.206954271 |
| imp*Lb*dCas12a | 12.17949 | 0.35222798 |

**Source Data Table 2. Source data of PAM-investigating fractions, used for the bar plot of Figure 3D. The other fractions, totalling to 1, are also reported.**

| | PAM-investigating | | 1D | | 3D | |
|---|---|---|---|---|---|---|
| Nuclease | Prob (%) | St. dev. | Prob (%) | St. dev. | Prob (%) | St. dev. |
| *Spy*dCas9 | 0.330709 | 0.03501 | 0.275591 | 0.030371 | 0.393701 | 0.036079 |
| *Sp*RYdCas9 | 0.338498 | 0.031611 | 0.300177 | 0.02838 | 0.361325 | 0.034974 |
| *Lb*dCas12a | 0.351324 | 0.016209 | 0.198574 | 0.009969 | 0.450102 | 0.018814 |
| imp*Lb*dCas12a | 0.469588 | 0.030267 | 0.141941 | 0.010746 | 0.38847 | 0.02197 |

**Source Data Table 3. Source data of average diffusion coefficients ($\mu m^2/s$), used for the scatter plots of Figure 4B.**

| | *Spy*dCas9 | | *Sp*RYdCas9 | | *Lb*dCas12a | | imp*Lb*dCas12a | |
|---|---|---|---|---|---|---|---|---|
| Track length | Avg. D* | St. dev. | Avg. D* | St. dev. | Avg. D* | St. dev. | Avg. D* | St. dev. |
| 3 | 0.57421921 | 0.006045 | 0.542011 | 0.00624 | 0.784748 | 0.008135 | 0.69277 | 0.007546 |
| 4 | 0.48265905 | 0.005354 | 0.4458916 | 0.005573 | 0.737751 | 0.007861 | 0.6554 | 0.00823 |
| 5 | 0.42657774 | 0.004586 | 0.3827837 | 0.004809 | 0.718802 | 0.007629 | 0.628498 | 0.008021 |
| 6 | 0.38228715 | 0.004047 | 0.3557168 | 0.004466 | 0.736711 | 0.007028 | 0.62063 | 0.007804 |
| 7 | 0.36854012 | 0.003393 | 0.3315628 | 0.004071 | 0.72226 | 0.005696 | 0.600112 | 0.007397 |
| >8 | 0.39834597 | 0.002726 | 0.333857 | 0.005361 | 0.734685 | 0.004565 | 0.630357 | 0.00624 |

**Source Data Table 4. Source data of average probability of 1 unit of each Cas nuclease to find 1 target inside the *E. coli* genome, used for scatter plot of Figure 5A (continuous lines).**

| | *Spy*dCas9 | | *Sp*RYdCas9 | | *Lb*dCas12a | | imp*Lb*dCas12a | |
|---|---|---|---|---|---|---|---|---|
| Time (min) | Avg. prob. | St. dev. | Avg. prob. | St. dev. | Avg. prob. | St. dev. | Avg. prob. | St. dev. |
| 1 | 0.33 | 0.1006 | 0.2 | 0.0943 | 2.065 | 0.1617 | 0.44 | 0.163 |
| 5 | 1.645 | 0.1536 | 0.935 | 0.231 | 10.68 | 0.4934 | 2.195 | 0.3647 |
| 10 | 3.15 | 0.3291 | 1.7 | 0.2896 | 19.575 | 0.5181 | 4.075 | 0.5361 |
| 30 | 9.76 | 0.8171 | 5.54 | 0.395 | 49.93 | 1.3907 | 11.94 | 0.6306 |
| 60 | 18.525 | 0.8545 | 10.39 | 0.6501 | 74.1 | 0.5788 | 21.62 | 0.68 |
| 120 | 33.015 | 1.1188 | 19.71 | 0.914 | 93.315 | 0.7835 | 39.115 | 0.886 |
| 240 | 55.15 | 1.0904 | 35.315 | 1.0483 | 100 | 0 | 63.755 | 0.7286 |
| 480 | 79.99 | 0.8736 | 59.04 | 0.7923 | 100 | 0 | 86.57 | 0.7436 |

**Source Data Table 5. Source data of average probability of 10 units of each Cas nuclease to find 1 target inside the *E. coli* genome, used for scatter plot of Figure 5A (dashed lines).**

| | ***Spy*dCas9** | | ***Sp*RYdCas9** | | ***Lb*dCas12a** | | **imp*Lb*dCas12a** | |
|---|---|---|---|---|---|---|---|---|
| **Time (min)** | **Avg. prob.** | **St. dev.** | **Avg. prob.** | **St. dev.** | **Avg. prob.** | **St. dev.** | **Avg. prob.** | **St. dev.** |
| 1 | 3.41 | 0.3044 | 1.69 | 0.2923 | 20.735 | 0.6876 | 4.155 | 0.376 |
| 5 | 15.45 | 0.7785 | 8.665 | 0.5391 | 68.29 | 1.0096 | 18.795 | 1.0114 |
| 10 | 28.675 | 1.1304 | 16.825 | 1.0191 | 89.49 | 0.6096 | 33.66 | 1.0456 |
| 30 | 63.54 | 0.667 | 42.65 | 1.2383 | 99.89 | 0.0394 | 71.685 | 0.6684 |
| 60 | 86.995 | 0.8821 | 67.02 | 0.7173 | 100 | 0 | 91.87 | 0.4015 |
| 120 | 98.175 | 0.3129 | 88.74 | 0.9101 | 100 | 0 | 99.315 | 0.2274 |
| 240 | 99.965 | 0.0412 | 98.95 | 0.2646 | 100 | 0 | 100 | 0 |
| 480 | 100 | 0 | 99.99 | 0.0211 | 100 | 0 | 100 | 0 |

**Source Data Table 6. Source data of $t_{50\%}$ expressed in minutes at varying GC content percentages, used for the scatter plot in figure 5B (top).**

| | **$t_{50\%}$ (min)** | | | |
|---|---|---|---|---|
| **GC content (%)** | ***Spy*dCas9** | ***Sp*RYdCas9** | ***Lb*dCas12a** | **imp*Lb*dCas12a** |
| 30.0 | 98 | 371 | 66 | 192 |
| 40.0 | 152 | 369 | 46 | 185 |
| 50.5 | 205 | 370 | 30 | 164 |
| 60.0 | 249 | 371 | 17 | 155 |
| 70.0 | 270 | 373 | 8.5 | 153 |

**Source Data Table 7. Source data of the number of PAM sequences at varying GC content percentage, used for the scatter plot in Figure 5B (bottom).**

| | **Number of PAM sequences** | | | |
|---|---|---|---|---|
| **GC content (%)** | ***Spy*dCas9** | ***Sp*RYdCas9** | ***Lb*dCas12a** | **imp*Lb*dCas12a** |
| 30.0 | 481,900.2 | 21,417,790 | 596,887 | 3,291,935 |
| 40.0 | 856,711.5 | 21,417,790 | 404,796.2 | 2,767,170 |
| 50.5 | 1,365,518 | 21,417,790 | 244,347 | 2,324,340 |
| 60.0 | 1,927,601 | 21,417,790 | 137,073.8 | 2038970 |
| 70.0 | 2,586,332 | 21,417,790 | 64,375.98 | 1,887,440 |

**Source Data Table 8. Source data of target-bound fractions at different number of tracks per cell, used for the scatter plot of Figure 6C.**

| | ***Spy*dCas9** | | ***Sp*RYdCas9** | | ***Lb*dCas12a** | | **imp*Lb*dCas12a** | |
|---|---|---|---|---|---|---|---|---|
| **Apparent copy number** | **Target-bound** | **St. dev.** | **Target-bound** | **St. dev.** | **Target-bound** | **St. dev.** | **Target-bound** | **St. dev.** |
| 7-50 | 15 | 4 | 5 | 2 | 35 | 4 | 29 | 5 |
| 51-100 | 13 | 4 | 3 | 1 | 30 | 2 | 26 | 3 |
| 101-150 | 10 | 2 | 3.6 | 1 | 28 | 4 | 20 | 3 |
| 151-200 | 3 | 2 | 3.2 | 2.5 | 25 | 3 | 18 | 3 |
